# Detecting cortical index of arousal in sleeping patients with unresponsive wakefulness syndrome

**DOI:** 10.1101/2025.08.02.668275

**Authors:** Man Li, Jiahui Pan, Junrong Han, Xiaoyu Bao, Hang Wu, Fei Wang, Yangzuyi Yu, Aoqi Li, Yuhao Liu, Di Chen, Xiaochun Yang, Yanbin He, Pengmin Qin, Yuanqing Li

## Abstract

Consciousness is assumed to be defined by two components: awareness and arousal. However, compared with awareness, the cortical index for arousal remains lacking in clues. Since patients with unresponsive wakefulness syndrome (UWS) are supposed to have no awareness but arousal, sleep-wake cycle in patients with UWS may reflect pure arousal fluctuations, and then could highlight the cortical indexes for arousal. This study recorded nighttime polysomnography in patients with UWS, patients with minimally conscious state and healthy controls. The EEG signal showed that spectral slope could index arousal, fluctuating among sleep stages in all three groups. Both spectral entropy and Lempel-Ziv complexity indexed awareness, showing a significant difference between conscious and unconscious states. These findings provide fundamental evidence for the two-component hypothesis of consciousness.

## INTRODUCTION

The debate on consciousness has attracted worldwide attention. In one popular theory, consciousness could be defined by two components: awareness (subjective experience, the content of consciousness) and arousal (the levels of wakefulness)^1–3^. Most states of consciousness can be described from these two components (Figure 1A). Different from awake healthy people who have high arousal and awareness, general anesthesia, N3-sleep and coma have low arousal and low awareness^2^. Rapid eye movement sleep (REM-sleep) has relatively high awareness but low arousal as the dreams commonly happen in REM-sleep and reflect rich subjective experience^4^. Patients with unresponsive wakefulness syndrome (UWS, also known as vegetative state) have no awareness but arousal as they have sleep-wake cycle, reflex response and high neural activity in brain stem ascending reticular activating system^5^. Although the majority of studies on consciousness (especially, disorders of consciousness) follow this two-component hypothesis of consciousness, compared with the plenty of studies on cortical indexes for awareness, little is known about the cortical indexes for arousal.

**Figure 1:**
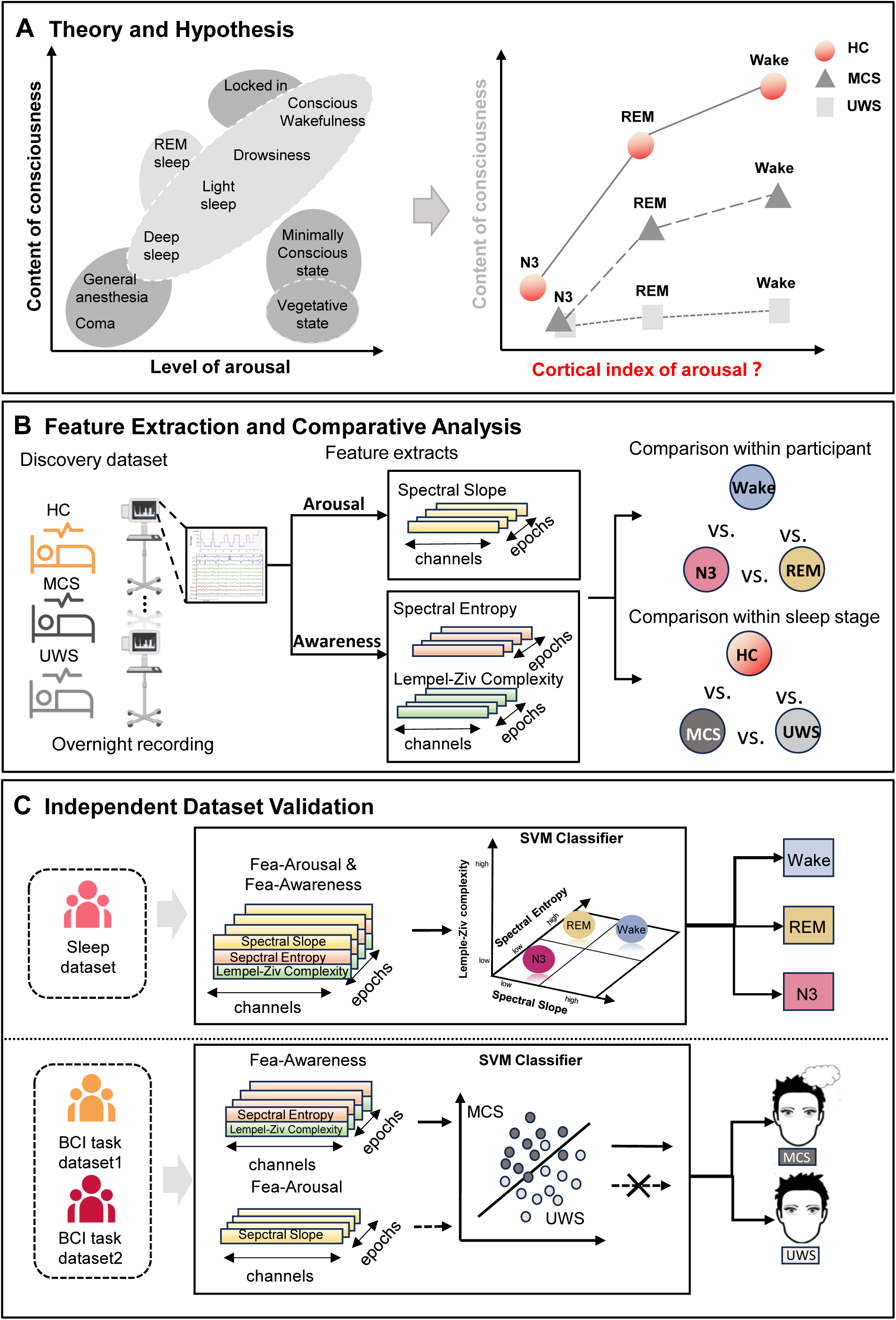
Overview of the study design. **(A)** Theory and Hypothesis. **(B)** Feature Extraction and Comparative Analysis: Detailing the EEG feature computation and the comparative analysis across different groups and sleep stages. **(C)** Independent Dataset Validation: Sleep staging and consciousness state diagnosis based on arousal and awareness-related features.

Different from cortical indexes for awareness which can be detected by comparing the wake states of healthy controls and UWS patients because both of them have high arousal, the cortical indexes for arousal are more difficult to investigate. As in most states of consciousness, arousal fluctuates along with awareness (Figure 1 A). For example, in sleep-wake cycle of healthy person, both arousal and awareness keep decreasing from awake state to N3-sleep^6^. Although REM-sleep was usually adopted to investigate the biomarker for awareness by comparing with N3-sleep, some previous studies suggested that REM-sleep also had higher arousal than N3-sleep^7–9^ (Figure 1A). Thus, the covariation of arousal and awareness generates huge obstacles for distinguishing the independent cortical indexes for arousal from ones for awareness, and even undermines the foundation of the two-component hypothesis of consciousness (there might not be independent arousal or awareness in consciousness at all). Facing this dilemma, sleep-wake cycle in patient with UWS provides a solution because patient with UWS is supposed to have no awareness, and then could present the pure arousal fluctuation among the sleep stages. Most importantly, this solution is feasible now. As recent studies had demonstrate that polysomnography (PSG) could successfully discriminate sleep stages in UWS^10–13^, and could acquire electroencephalography (EEG) signals simultaneously which has been commonly used in the investigations on neural substrate of consciousness^10,14–18^.

What measurements on EEG signals could be candidates for cortical indexes for arousal? The majority of previous EEG studies on consciousness focused on cortical indexes for awareness. They found that the spectral entropy and Lempel-Ziv complexity of EEG signal, reflecting the richness and diversity of neural dynamics, could be regarded as the markers of awareness^19,20^. The Perturbational Complexity Index, combining TMS-EEG and Lempel-Ziv complexity, has been used to distinguish UWS from minimally conscious state (MCS) who have weak and unstable consciousness^19,21^. Additionally, alpha power is also a usually mentioned EEG measurement which was suggested to be related to levels of awareness, showing significant reduction in UWS in some previous studies^14,20,22,23^. Different from the measurements of EEG complexities, another widely concerned measurement is the spectral slope of the EEG power spectrum. It represents the scale-free component of cortical activity, could reflect the levels of consciousness according to the results from sleep and anesthesia states^24–28^. As the above mentioned, the sleep and anesthesia showed the both reduction of awareness and arousal, these finding might suggest spectral slope also had the probability to reflect the levels of arousal^29^. Based on these EEG measurements in the previous studies, we assumed that it should be achievable to disclose the distinct cortical indexes for arousal by adopting the sleep-wake cycle in both healthy participants and UWS participants.

To achieve the above aim, the current study included 67 UWS, 55 MCS and 55 healthy controls (HC) in total. See Table 1 for detailed information. The whole study included two parts (Figure 1). In the first part, PSG was used to record nighttime EEG, along with other physiological signals (EOG and EMG) in 16 UWS, 18 MCS and 18 HC. The EEG signals within Wake, N3-sleep, and REM-sleep stages were used to compute spectral slope, spectral entropy, and Lempel-Ziv complexity for each group (Figure 1B). Secondly, the findings from the first part were further validated using an independent sleep EEG dataset in healthy participants (n=37) and two independent BCI task-related EEG datasets (the first including 45 UWS, 33 MCS, the second including 6 UWS, 4 MCS) (Figure 1C). Following two-components hypothesis of consciousness, UWS would have lower awareness than HC and MCS during both Wake state and REM-sleep, but have the similar arousal as the HC and MCS among Wake state, REM-sleep and N3-sleep (Figure 1A). Our results showed that the spectral slope can work as cortical index for arousal, distinct from both the spectral entropy and Lempel-Ziv complexity which work as the indexes for awareness. These findings support the two-component theory of consciousness, and shed light on the biological substrate of consciousness.

**Table 1:** Dataset Information.

| Dataset | State | Subjects | Gender (M/F) | Age (Mean ± SD) | ABI (%) | TBI (%) | CVD (%) |
| --- | --- | --- | --- | --- | --- | --- | --- |
| Sleep dataset1 | UWS | 16 | 9/7 | 51.68 ± 11.05 | 31.25% | 37.50% | 31.25% |
| | MCS | 18 | 15/3 | 51.27 ± 9.63 | 5.55% | 44.44% | 50.00% |
| | HC | 18 | 8/10 | 41.0 ± 9.99 | — | — | — |
| Sleep dataset2 | HC | 37 | 20/17 | 26.45 ± 8.11 | — | — | — |
| BCI task dataset1 | UWS | 45 | 31/14 | 39.06 ± 14.17 | 26.66% | 42.22% | 31.11% |
| | MCS | 33 | 20/13 | 36.24 ± 14.48 | 15.15% | 66.67% | 18.18% |
| BCI task dataset2 | UWS | 6 | 4/2 | 49.17 ± 8.45 | 50.00% | 50.00% | 0.00% |
| | MCS | 4 | 3/1 | 51.75 ± 19.35 | 75.00% | 25.00% | 0.00% |

**Note.**Dataset specifies the data collection used in this study, including two Sleep datasets and two BCI task datasets. State represents the clinical state of the participants (UWS, MCS, or HC). Gender is reported as male/female, and Age as mean ± standard deviation. ABI, TBI, and CVD indicate the percentages of participants diagnosed with anoxic brain injury, traumatic brain injury, and combined cerebrovascular/inflammatory diseases, respectively. “—” denotes unavailable data. The demographic information for each dataset is provided in the Supplemental Information.

## RESULTS

### Altered Sleep Pattern in UWS

The sleep pattern of patients with UWS appeared to be characterized by frequent changes between sleep and wakefulness^30^. To evaluate group differences, several sleep architecture metrics including total sleep time (TST), wake time after sleep onset (WASO), sleep efficiency (SE), sleep maintenance efficiency (SME), night wake dominance (NWD), and sleep architecture completeness (SAC) were quantified. Figure 2 shows expert-scored hypnograms for the HC, MCS, and UWS groups.

**Figure 2:**
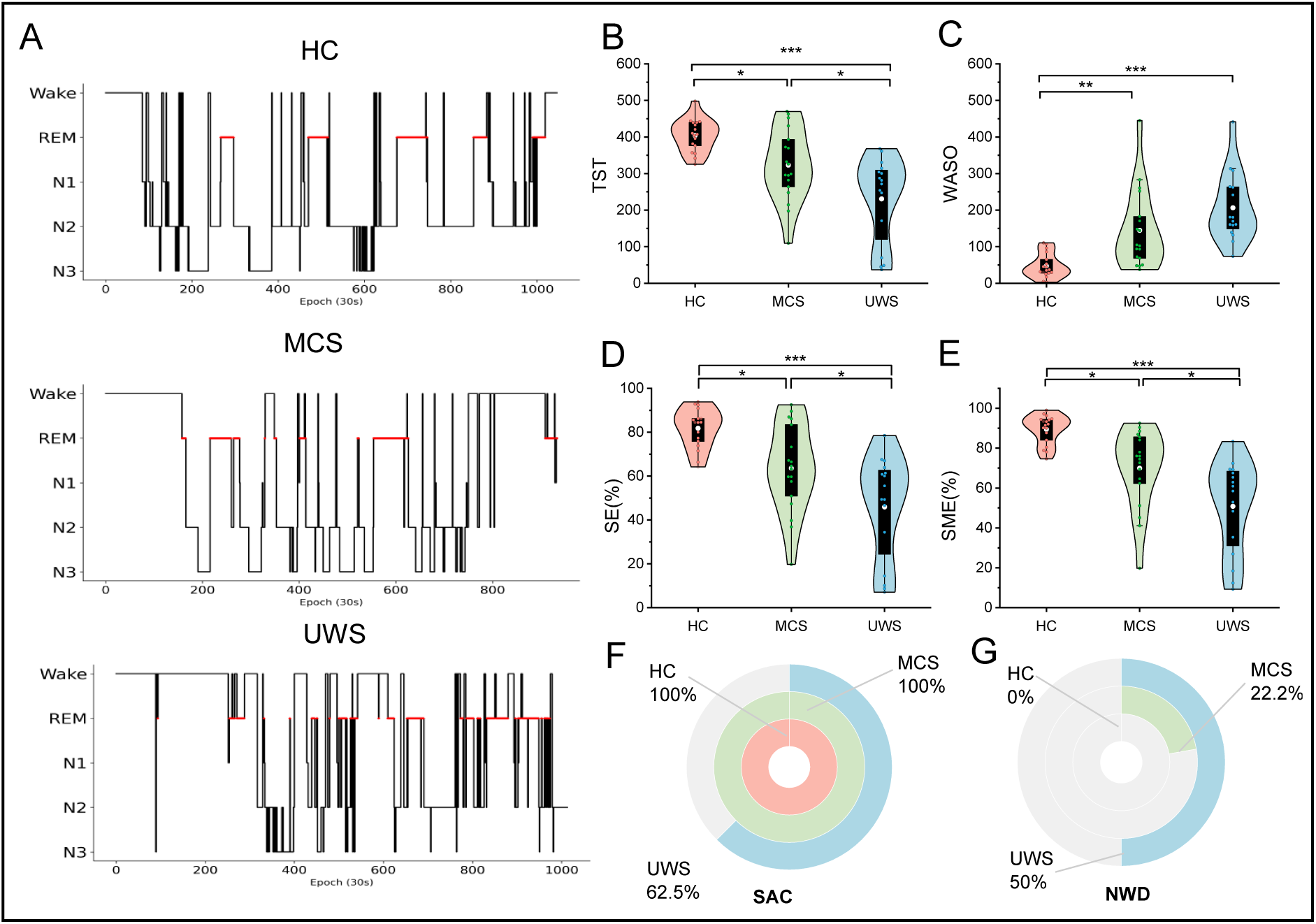
Sleep Patterns. **(A)** Exemplary hypnograms: Upper panel represents healthy controls (HC), middle panel depicts minimally conscious state (MCS), and lower panel shows unresponsive wakefulness syndrome (UWS). **(B)** Total sleep time (TST) for each group, defined as the cumulative duration of N1, N2, N3, and REM sleep stages within the Sleep Period Time (SPT; interval from first to last sleep epoch). **(C)** Wake after sleep onset (WASO) for each group, representing total wake duration within SPT. **(D)** Sleep efficiency (SE), calculated as TST / Time in Bed (TIB) × 100 (%), where TIB reflects the total recording duration. **(E)** Sleep maintenance efficiency (SME), calculated as TST / SPT × 100 (%). **(F)** Sleep architecture completeness (SAC): Percentage of participants in each group exhibiting all standard sleep stages (N1, N2, N3, and REM) during the recorded sleep period. **(G)** Night wake dominance (NWD): Percentage of participants with greater wake duration than sleep time during nocturnal hours. The white dot indicates the mean, and the box boundaries indicate the upper and lower quartiles. ∗*p* < 0.05, ∗ ∗ *p* < 0.01, ∗ ∗ ∗*p* < 0.001.

One-way ANCOVAs were performed to compare these standard sleep metrics from hypnogram between groups (HC, MCS, UWS), respectively, with age as a covariate to control for potential age-related effects. Post-hoc pairwise comparisons with Bonferroni correction were conducted to assess differences between groups. All the sleep metrics showed significant differences among groups (TST: *F* (2, 47) = 13.971, *p <* 0.001, *η*^2^ = 0.373; WASO: *F* (2, 48) = 14.401, *p <* 0.001, *η*^2^ = 0.375; SE: *F* (2, 47) = 13.728, *p <* 0.001,*η*^2^ = 0.369; SME: *F* (2, 48) = 17.104, *p <* 0.001, *η*^2^ = 0.416). TST was significantly higher in the HC group compared with the MCS (*p* = 0.03) and UWS groups (*p <* 0.001). MCS group also exhibited higher values than the UWS group (*p* = 0.016). For WASO, UWS and MCS patients exhibited more frequent sleep-wake transitions than HC group (UWS vs. HC: *p <* 0.001; MCS vs. HC: *p* = 0.003). For SE, HC group had higher values than both the MCS (*p* = 0.022) and UWS groups (*p <* 0.001), as well as the MCS group had higher value than the UWS group (p=0.025). Similarly, for SME, the HC group had significantly higher values compared to the MCS (*p* = 0.01) and UWS groups (*p <* 0.001), with the MCS group also exhibiting higher values than the UWS group (*p* = 0.01) (Figure 2B-E). Please see Supplemental Information Table S1 for detailed statistical information. Notable differences were observed in SAC and NWD among the three groups, with the UWS group exhibiting the lowest SAC and the most pronounced NWD. As illustrated in Figure 2F-G, only 62.5% of UWS patients exhibited the presence of all standard sleep stages. Furthermore, NWD was absent in HC participants but present in 22.2% of MCS patients and 50% of UWS patients. These results highlight the distinct sleep architecture of each group, characterized by impaired sleep efficiency and maintenance in UWS patients, in contrast to the more stable sleep patterns observed in the HC and MCS groups.

### Spectral slope as the cortical index for Arousal

Among the EEG measurements mentioned above, only the spectral slope fits the hypothesis on the arousal. The spectral slope reliably tracked arousal variations across sleep stages in all three groups, especially in UWS. Additionally, the spectral slope in HC shows no difference with MCS and UWS during Wake state. See the following for the detailed information.

For the HC and MCS groups, a one-way repeated measures ANOVA was used to assess the effects of sleep stage. The results showed significant differences in spectral slope between sleep stages within each group (HC: *F* (1.404, 23.860) = 242.340, *p <* 0.001, *η*^2^ = 0.934; MCS: *F* (2, 34) = 27.661, *p <* 0.001, *η*^2^ = 0.619). In post-hoc comparisons with Bonferroni correction, the HC group showed significantly higher spectral slope during Wake state compared to REM-sleep (*p <* 0.001) and N3-sleep (*p <* 0.001), as well as significantly higher spectral slope during REM compared to N3 (*p <* 0.001). Similarly, in the MCS group, spectral slope was highest in Wake state, followed by REM-sleep and N3-sleep, with significant differences between Wake and REM (*p <* 0.001), Wake and N3 (*p <* 0.001), and REM and N3 (*p* = 0.011). For the UWS group, due to missing data for certain sleep stages in several patients (Supplemental Information, Table S2), a one-way ANCOVA was performed for comparison between sleep stages, with age included as a covariate. The results showed significant difference among Wake, REM-sleep and N3-sleep (*F* (2, 37) = 9.259, *p <* 0.001, *η*^2^ = 0.334). Post-hoc analysis with Bonferroni correction showed that the spectral slope was also significantly higher during Wake compared to REM-sleep (*p* = 0.039) and N3-sleep (*p <* 0.001) (Figure 3A).

**Figure 3:**
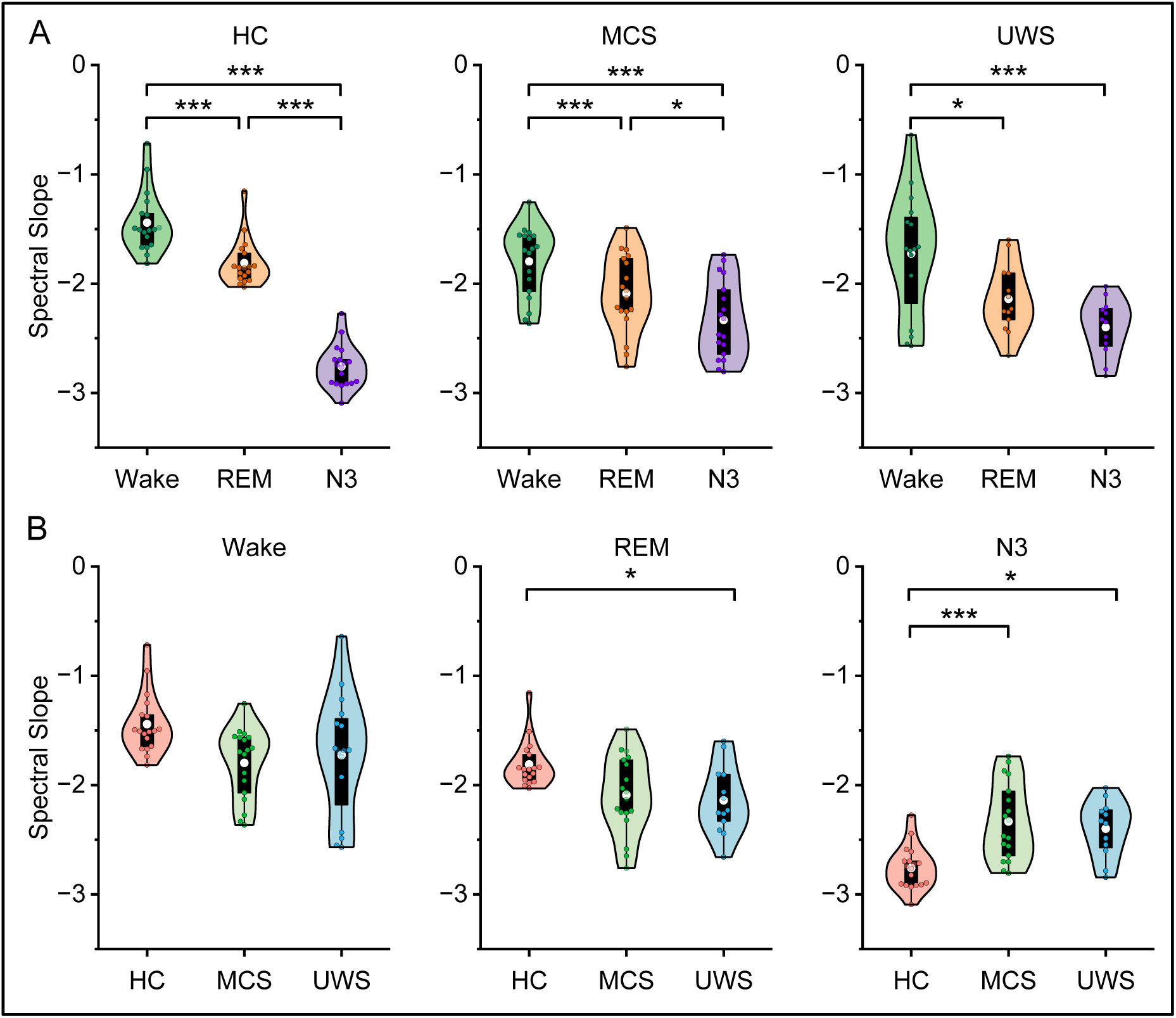
Arousal-Related Spectral Slope Trends. **(A)** Violin plots of spectral slope distributions for each sleep stage within HC, MCS, and UWS groups. **(B)** Violin plots of spectral slope distributions for individual sleep stages across HC, MCS, and UWS groups. The white dot indicates the mean, and the box boundaries represent the upper and lower quartiles.∗*p <* 0.05, ∗ ∗ *p <* 0.01, ∗ ∗ ∗*p <* 0.001.

Furthermore, one-way ANCOVA was performed to compare spectral slope between groups (HC, MCS, UWS) for each sleep stage with age as a covariate, respectively. There were no significant differences among groups in Wake stage, while there were significant differences between among in REM and N3 stage (REM: *F* (2, 45) = 4.02, *p* = 0.025, *η*^2^ = 0.152; N3: *F* (2, 44) = 8.271, *p <* 0.001, *η*^2^ = 0.273). For post-hoc comparisons with Bonferroni corrections, during REM stage, spectral slope in HC group was significantly higher than that in the UWS group (*p* = 0.04), a marginally significant differences were observed between HC and MCS (*p* = 0.059), while no difference between MCS and UWS. During N3-sleep, HC group exhibited significantly lower spectral slope compared to both MCS (*p <* 0.001) and UWS (*p* = 0.014), no significant difference was found between MCS and UWS (Figure 3B). Please see Supplemental Information Tables S3-5 for detailed statistical information.

To further examine the influence of etiology, we compared spectral slope between anoxia and no-anoxia subgroups within MCS and UWS patients (MCS: 1/17; UWS: 5/11). In the UWS anoxia subgroup, a Kruskal–Wallis test revealed a significant effect of sleep stage (*p* = 0.033). Post-hoc comparisons indicated that spectral slope was higher during Wake compared to N3 (*p* = 0.039). In contrast, the UWS no-anoxia subgroup showed no significant stage-related differences (*p* = 0.08). In the MCS no-anoxia subgroup, spectral slope also differed significantly across sleep stages as shown by repeated-measures ANOVA (*F* (2, 32) = 24.631, *p <* 0.001,*η*^2^ = 0.606). Post-hoc tests with Bonferroni correction indicated that, spectral slope was higher during Wake compared to REM (*p* = 0.02) and N3 (*p <* 0.001), and during REM compared to N3 (*p* = 0.012) (Supplemental Information Figure S1A). Finally, between-group comparisons using Mann–Whitney U tests revealed no significant differences between MCS and UWS under no-anoxia conditions across Wake, REM, and N3 (Supplemental Information Figure S1B). These results demonstrated that the spectral slope primarily reflects the arousal dimension of consciousness, rather than the awareness dimension.

### The measurements of complexity as cortical index for Awareness

Different from spectral slope, the measurements of complexity (spectral entropy and Lempel-Ziv complexity) fit the hypothesis on awareness. HC and MCS showed higher complexity during Wake state and REM-sleep than during N3-sleep, while UWS showed no fluctuation among the sleep stages. Additionally, HC and MCS showed higher complexity than UWS in both Wake state and REM-sleep.

Figure 4A illustrates the variations in spectral entropy across the sleep stages within each group. One-way repeated-measures ANOVA revealed significant differences in spectral entropy across sleep stages in the HC and MCS groups (HC: *F* (1.283, 21.803) = 66.073, *p <* 0.001, *η_p_*^2^ = 0.795; MCS: *F* (1.490, 25.328) = 21.773, *p <* 0.001, *η_p_*^2^ = 0.562). Post-hoc comparisons with Bonferroni correction showed that, in the HC group, spectral entropy was significantly higher during Wake state compared to N3-sleep (*p <* 0.001), higher during REM-sleep compared to N3-sleep (*p <* 0.001). In the MCS group, post-hoc tests with Bonferroni correction revealed significant higher spectral entropy being higher in Wake state compared to N3-sleep (*p <* 0.001), as well as in REM-sleep compared to N3-sleep (*p <* 0.001). Most importantly, in the UWS group, no significant differences were found among sleep stages, indicating the absence of awareness modulation among the wake-sleep cycle while the arousal still fluctuated independently.

**Figure 4:**
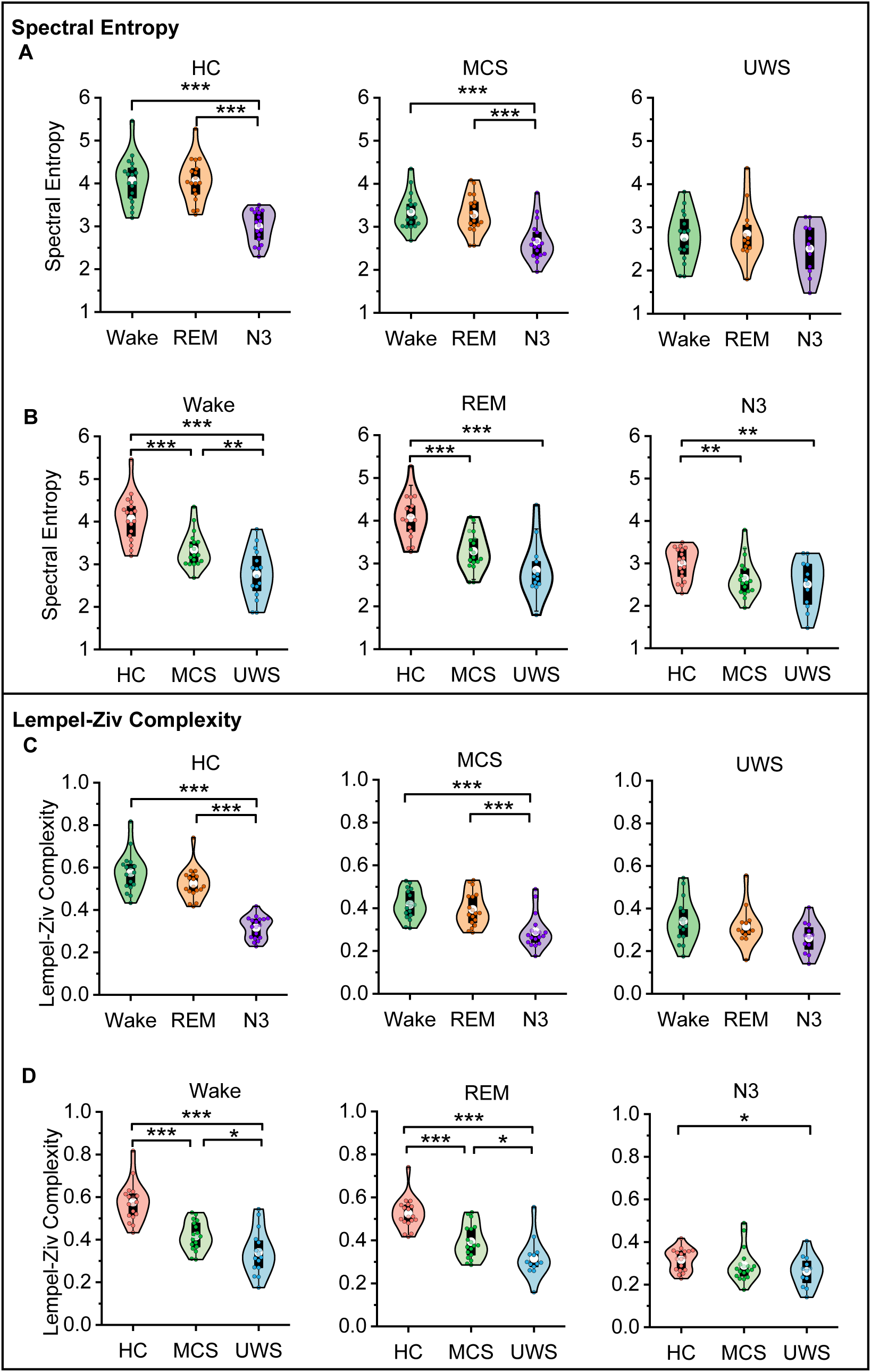
Spectral Entropy and Lempel-Ziv Complexity Trends. **(A)**Mean spectral entropy variations across Wake, N3, and REM stages, combining data from HC, MCS, and UWS groups. **(B)** Violin plots of spectral entropy distributions for each sleep stage within HC, MCS, and UWS groups. **(C)** Mean Lempel-Ziv complexity variations across Wake, N3, and REM stages, combining data from HC, MCS, and UWS groups. **(D)** Violin plots of Lempel-Ziv complexity distributions for individual sleep stages across HC, MCS, and UWS groups. The white dot indicates the mean, and the box boundaries represent the upper and lower quartiles.∗*p <* 0.05, ∗ ∗ *p <* 0.01, ∗ ∗ ∗*p <* 0.001.

To further validate the role of spectral entropy as a marker of awareness, one-way ANCOVA was performed to compare spectral entropy between groups (HC, MCS, UWS) for each sleep stage, with age as a covariate to control for potential age-related effects (Figure 4B). The results of one-way ANCOVA showed significant difference between groups in all Wake state, REM-sleep and N3-sleep (Wake state: *F* (2, 48) = 26.139, *p <* 0.001, *η_p_*^2^ = 0.521; REM-sleep: *F* (2, 45) = 22.422, *p <* 0.001, *η_p_*^2^ = 0.499; N3-sleep: *F* (2, 44) = 8.008, *p* = 0.001, *η_p_*^2^ = 0.267). In post-hoc comparisons with Bonferroni correction, in the Wake state, spectral entropy was significantly higher in the HC group compared to both the MCS (*p <* 0.001) and UWS groups (*p <* 0.001), with the MCS group also exhibiting significantly higher spectral entropy than the UWS group (*p* = 0.006). In the REM-sleep and N3-sleep stages, the HC group exhibited significantly higher spectral entropy compared to the MCS group (REM-sleep: HC vs. MCS, *p <* 0.001; N3-sleep: HC vs. MCS, *p* = 0.008) and UWS group (REM: HC vs. UWS, *p <* 0.001, N3: HC vs. UWS, *p* = 0.002). No significant differences were found between the MCS and UWS groups.

Lempel-Ziv complexity as another popular measurement for complexity, a one-way repeated measures ANOVA show significant differences of Lempel-Ziv complexity between sleep stages in HC and MCS group (HC: *F* (1.455, 24.743) = 127.414, *p <* 0.001, *η_p_*^2^ = 0.882; MCS: *F* (2, 34) = 27.550, *p <* 0.001, *η_p_*^2^ = 0.618). As illustrated in Figure 4C, post-hoc comparisons with Bonferroni-correction showed that, in the HC group, both Wake and REM-sleep showed higher Lempel-Ziv complexity than in N3-sleep (for Wake vs. N3-sleep: *p <* 0.001; for REM-sleep vs. N3-sleep: *p <* 0.001). In the MCS group, the post-hoc comparisons with Bonferroni-correction also showed that both Wake and REM-sleep showed higher Lempel-Ziv complexity than in N3-sleep (for Wake vs. N3-sleep: *p <* 0.001; for REM-sleep vs. N3-sleep: *p <* 0.001). However, in the UWS group, one-way ANCOVA (with age as a covariate) showed no significant differences were observed across sleep stages, indicating the absence of awareness modulation during sleep-wake cycle. As illustrated in Figure 4D, the result of one-way ANCOVA (with age included as a covariate) showed significant difference between groups in Wake, REM and N3 stage (Wake: *F* (2, 48) = 27.608, *p <* 0.001, *η_p_*^2^ = 0.535; REM: *F* (2, 45) = 25.246, *p <* 0.001, *η_p_*^2^ = 0.529; N3: *F* (2, 44) = 4.154, *p* = 0.022, *η*^2^ = 0.159). For post-hoc comparisons with Bonferroni-correction, during Wake and REM, Lempel-Ziv complexity in the HC group was significantly higher than in both MCS (Wake: *p <* 0.001; REM: *p <* 0.001) and UWS (Wake: *p <* 0.001; REM: *p <* 0.001), as well as Lempel-Ziv complexity in the MCS group was significantly higher than in the UWS group (Wake: *p* = 0.045; REM: *p* = 0.026). In the N3 stage, only the HC group exhibited significantly higher Lempel-Ziv complexity compared to the UWS group (*p* = 0.021). Please see Supplemental Information Tables S3-5 for detailed statistical information.

### Sleep Staging based on cortical index of arousal and awareness

To confirm the effectiveness of arousal and awareness-related features, we evaluated their contributions in sleep stage classification using three analytical approaches on an independent sleep dataset in healthy participants. Figure 5A presents the confusion matrices from the leave-one-subject-out cross-validation (LOSO-CV) analysis, illustrating three classification performances: combining both arousal and awareness-related features, using arousal-related features, and using awareness-related features. Specifically, the combined feature set achieved the highest overall accuracy (*ACC* = 0.80), macro-average F1 score (*MF* 1 = 0.7745), and Cohen’s Kappa (*K* = 0.6864). In contrast, using only arousal-related features resulted in the lowest classification performance (*ACC* = 0.6481, *MF* 1 = 0.6481, *K* = 0.4695), while the awareness-only feature set achieved intermediate performance (*ACC* = 0.7617, *MF* 1 = 0.7344, *K* = 0.6316).

**Figure 5:**
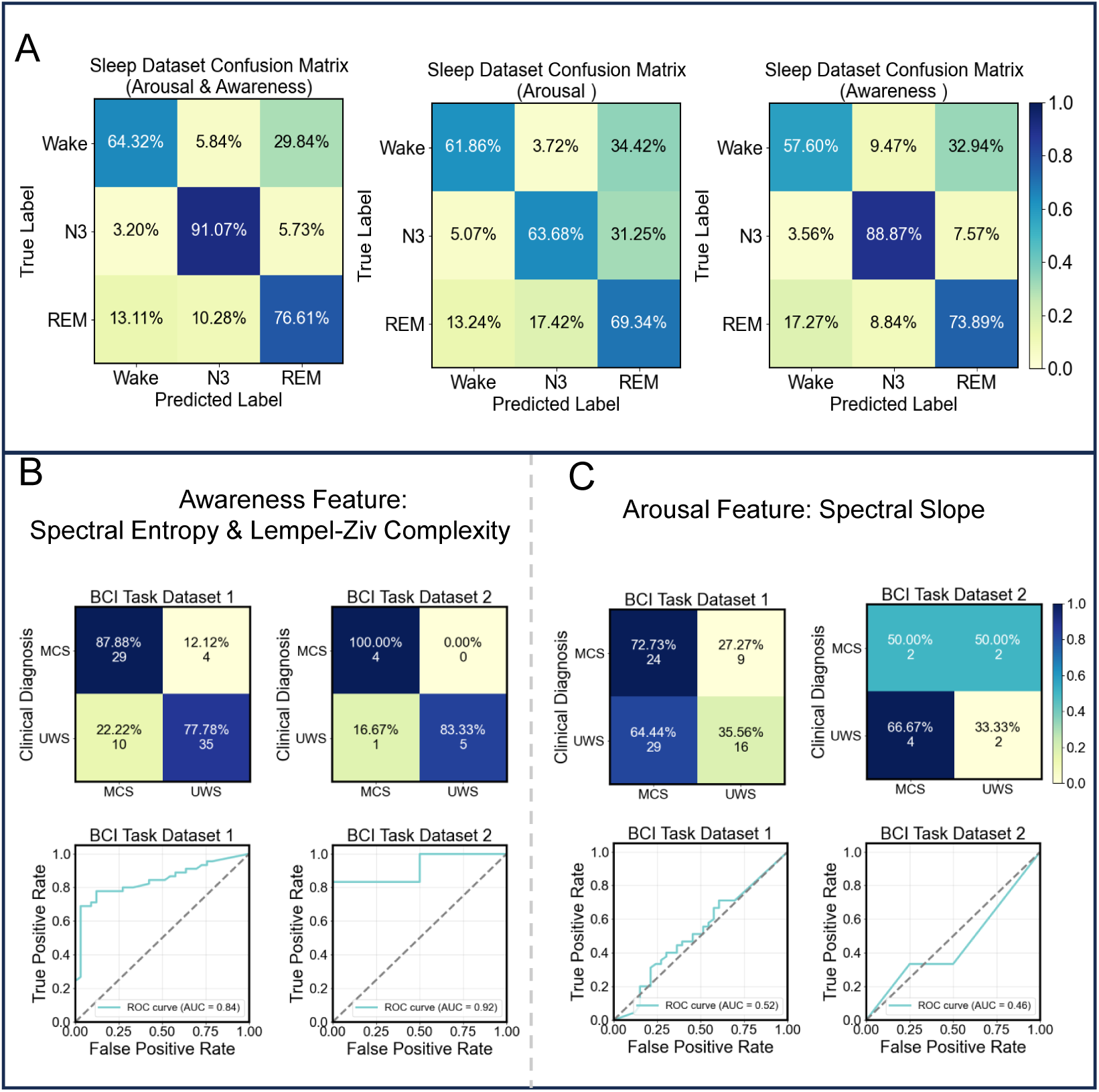
Evaluation of Sleep Staging Performance and Consciousness Diagnosis. **(A)** Cross-validated classification performance under different feature combinations. Left: Joint use of arousal-related (spectral slope) and awareness-related features (spectral entropy, Lempel-Ziv complexity) with leave-one-subject-out cross-validation (LOSO-CV). Middle: Arousal-related features (spectral slope). Right: Awareness-related features (spectral entropy, Lempel-Ziv complexity). **(B)**Awareness driven diagnostic performance: Top row, Confusion matrices for BCI task dataset1 (left) and BCI task dataset2 (right); Bottom row, ROC curves (BCI task dataset1 AUC: 0.84, BCI task dataset2 AUC: 0.92) confirm robust MCS-UWS spectral entropy arability. **(C)** Arousal driven diagnostic performance: Top row, Confusion matrices for identical datasets; Bottom row, ROC curves (dataset1 AUC: 0.52, dataset2 AUC: 0.46) approaching chance-level discrimination.

### Detecting consciousness based on cortical index of awareness

To further confirm the relationship between cortical indexes (awareness and arousal) and consciousness, we conducted classification analyses across two independent DOC groups (both including UWS and MCS). Both groups had the EEG datasets during brain-computer interface tasks (see Materials and Methods, Participants and Data Acquisition for details). The awareness-related features (spectral entropy and Lempel-Ziv complexity) achieved significantly higher classification performance (on UWS and MCS) compared to arousal-related features (spectral slope) which had no ability to discriminate UWS from MCS (see Figure 5B-C). Specifically, for the BCI task dataset1, the accuracy for awareness-related features was 82.05%, with an AUC of 0.84; and for the BCI task dataset2, the accuracy was 90%, with an AUC of 0.92. In contrast, arousal-related feature (spectral slope) resulted in much worse performance, with an accuracy of 51.28% and an AUC of 0.52 on the BCI task dataset1, and 40% accuracy with an AUC of 0.46 on the BCI task dataset2. These findings validate that spectral entropy and Lempel-Ziv complexity could index awareness while spectral slope, as the index of arousal, cannot index awareness.

### Reduced alpha power in healthy controls’ REM-sleep and UWS

For the HC and MCS groups, a one-way repeated measures ANOVA was used to assess the effects of sleep stage. The results showed significant differences in alpha power across sleep stages within each group (HC: *F* (1.458, 24.789) = 20.006, *p <* 0.001, *η_p_*^2^ = 0.541, MCS: *F* (1.372) = 9.420, *p* = 0.003, *η_p_*^2^ = 0.357). Post-hoc comparisons with Bonferroni correction showed that in the HC group, alpha power during REM-sleep was significantly lower than during Wake state (*p <* 0.001) and N3-sleep (*p <* 0.001). In the MCS group, both REM-sleep and N3-sleep exhibited significantly lower alpha power compared to Wake state (REM vs. Wake: *p <* 0.001; N3 vs. Wake: *p* = 0.017). However, in the UWS group, a one-way ANCOVA (with age as a covariate) revealed no significant differences in alpha power across sleep stages.

Furthermore, one-way ANCOVA was performed to compare alpha power between groups (HC, MCS, UWS) for each sleep stage with age as a covariate, respectively. The result showed significant difference between groups in Wake state, REM-sleep and N3-sleep (Wake: *F* (2, 48) = 26.848, *p <* 0.001, *η_p_*^2^ = 0.528; REM: *F* (2, 45) = 17.463, *p <* 0.001, *η_p_*^2^ = 0.437; N3: *F* (2, 44) = 20.700, *p <* 0.001, *η*^2^ = 0.485). Post-hoc comparisons with Bonferroni correction showed that alpha power in the UWS group was significantly lower than in both the HC and MCS groups during Wake (vs. HC: *p <* 0.001; vs. MCS: *p* = 0.003), REM (vs. HC: *p <* 0.001; vs. MCS: *p <* 0.001). In addition, the MCS group exhibited significantly lower alpha power than the HC group during Wake (*p <* 0.001) and N3 (*p <* 0.001). For detailed statistical results, see Supplemental Information Tables S3–S5 and Figure S2.

## DISCUSSION

The current study distinguishes the awareness from arousal with independent cortical indexes for each other. In specific, the spectral slope can work as a cortical index of arousal while the measurements of complexity (spectral entropy and Lempel-Ziv complexity) work as the cortical indexes of awareness. It is the first time to use cortical index describes the pure arousal fluctuation during sleep-wake cycle in patients with UWS, and demonstrates the hypothesis that the UWS has the similar arousal level as the healthy control. These findings strongly support the two-components hypothesis of consciousness.

The spectral slope works as an index for arousal. In the last two decades, most studies took the opinion that the patients with UWS have similar arousal level as healthy participants during wakefulness^31,32^. However, the evidence supporting this assumption is only from sleep-wake cycle in behavioral level^5,33,34^ and intact brain stem activity shown by PET^3,35^ in patients with UWS. Since there was no any evidence from cortical activity, these findings cannot eliminate the suspicion whether the patients with UWS have the similar brain arousal level as healthy individuals or not. In order to discover the cortical index for arousal, recent studies adopted fMRI or EEG, and acquired the signals from multiple altered conscious states (i.e., N3-sleep, deep anesthesia, REM-sleep and UWS)^26,29,32^. Unfortunately, in these studies, all the contrasts based on these states cannot rule out the mutual influence of the two components. For example, although REM-sleep have relative high awareness and low arousal^19,29^, it still shows higher value in both arousal and awareness compared to the N3-sleep. In the current study, following the hypothesis that patients with UWS have no awareness, their sleep-wake cycle could reflect the fluctuation of arousal purely. Consistent within this hypothesis, our results showed the spectral slope can reflect the arousal fluctuation among the sleep stages.

The spectral slope reflects a 1/f dynamic in EEG signals, i.e. higher frequency activity exhibits reduced power compared with lower frequency activity, which has been highlighted its importance in consciousness^36–38^. Previous studies indicated that the general anesthesia had more negative spectral slope values compared with awake states^24,39^. In the literature on sleep, within a brand frequency band (0.5-45Hz), N3-sleep showed the most negative spectral slope values compared with REM-sleep and Wake state while the REM-sleep had more negative spectral slope values than Wake state^27,40–42^. The current study showed consistent results with the previous studies in healthy participants. The most important finding of the current study was that fluctuations of the spectral slope across sleep stages in UWS patients were similar to those in healthy participants. As well as, spectral slope showed no difference among UWS, MCS, HC during Wake state, and could not discriminate the MCS from UWS in the two independent datasets. Although the previous studies indicated that the effects of anoxia on EEG signal in patients with UWS and MCS^43^, the current study showed that anoxia UWS patients also had more negative spectral slope during N3-sleep than Wake state, similar as the non-anoxia UWS patients. These results on spectral slope was in line with the assumption on cortical index of arousal.

The complexity of cortical activity indexes awareness. The links between complexity and awareness have been the focus of attention in last decade^44^. According to the integrated information theory (IIT) and similar ones^45,46^, a minimal level of complexity of cortical activities is needed to support conscious experience (e.g., awareness). In fact, some previous studies reported that complexity of cortical activities diminishes in the absence of awareness^42,47–50^. The spectral entropy and Lempel-Ziv complexity, two popular measurements of complexity, showed reduced values in unconscious states (UWS, N3-sleep, and anesthesia) compared with conscious states (awake, REM-sleep)^19,22,51–54^. Beyond these previous studies, our results showed that spectral entropy and Lempel-Ziv complexity did not fluctuate with sleep-wake cycle in UWS where the arousal (e.g. spectral slope) did, and could discriminate MCS from UWS patients in the two independent datasets. These further clarify the relationship between complexity and awareness. In current study, although most conscious states showed significant higher spectral entropy values than unconscious states, MCS patients had higher spectral entropy value (but not significant) than UWS patients during REM-sleep. This indicates that a single measurement may be unconvincing in a few cases, and combining multiple measurements will be more effective to discriminate MCS from UWS in clinic. Finally, both spectral entropy and Lempel-Ziv complexity showed higher value in healthy controls than UWS during N3-sleep. This may indicate that there is more information processing in unconscious healthy brain than UWS, which is supported by the previous studies on the cognitive processing during N3-sleep^55–57^.

Both UWS and MCS had detectable sleep stages during night-time. However, they showed altered sleep structure compared with healthy controls. Compared with MCS patients, UWS patients had shorter sleep duration. Additionally, there were 62.5% of the UWS patients had all sleep stages while all MCS patients had all sleep stages (Figure 2, Supplemental Information Table S2). These results were consistent with the previous studies^13,30,34,58–62^. Besides the difference of sleep structure between the levels of consciousness (UWS and MCS), the previous studies also intended to investigate the relationship between sleep elements and the outcome of the patients with UWS and MCS^10,63–67^, and found the positive relationship between sleep structure and better outcome^11,68,69^. For example, the recent studies showed that the sleep spindles, K-complex, and REM-sleep could predict the favorable outcome of patients^10,66,67^. The well-formed sleep spindles could even predict the detection of patients with cognitive-motor dissociation (CMD)^10^. All the above findings indicated that the sleep structure could reflect the remaining brain function and have important clinical values in patients with UWS and MCS. This study extended the current knowledge, for the first time describing the pure arousal fluctuation during sleep-wake cycle using cortical index (e.g. spectral slope).

The relationship between alpha power and awareness was also tested as some previous studies suggested that alpha power could be a biomarker of awareness^14,20,22,23,70–73^. Consistent with the studies, our results showed that healthy controls and MCS had stronger alpha power than UWS during Wake state. Furthermore, the current study showed that there was no difference among Wake state, REM-sleep and N3-sleep in UWS. However, our results also showed that both Wake state and N3-sleep had stronger alpha power than REM-sleep in healthy controls (see Supplemental Information Figure S2 and S3), which was also supported by previous study^74^. Considering that one in REM-sleep might have more conscious experience (e.g., awareness) than N3-sleep, this result might indicate that alpha could not reflect the level of awareness. Since the locked-in syndrome, who had lesions in brain stem, also showed reduced alpha power during awake states^75^, the reduced alpha power in REM-sleep might due to the reduced functional integrity between brain stem and cortices^75,76^. Furthermore, one recent study found that alpha power was suppressed only in UWS with severe post-anoxia but failed to discriminate un/consciousness in other aetiologies^43^. They supposed that the alpha power reduction might be related to the diffuse cortical damage. Taken together, the alpha power might not index awareness.

Several issues need to be noticed. First, although the most previous studies indicated that the REM-sleep might have higher arousal than N3-sleep as it has more brain activities^7,8^, some other studies showed REM-sleep had more negative spectral slope values compared with NREM-sleep and wake state within a narrow frequency band (30-45Hz or 30-50Hz)^27,29^. The current study reproduced this finding in healthy participants but not in UWS (see Supplemental Information Figure S4 and S5). This indicated that the narrow band spectral slope might not work as a biomarker of arousal. Second, N3-sleep showed more reduced spectral slope in healthy controls compared with UWS and MCS, which may indicate the intact function of brainstem and basal forebrain^65^. Finally, a few studies indicated that UWS patients showed more negative spectral slope than MCS during resting awake awake^43^. One possible reason was that the UWS patients might have more transition from awake state to sleep during resting states, resulting in a lower spectral slope compared with MCS patients. In this study, we used the EEG data from patients performing active tasks to minimize the influence of altered levels of consciousness. The results showed that spectral slope failed to discriminate between UWS and MCS, indicating the spectral slope cannot index awareness (Figure 5C).

In conclusion, combining the brain signal during sleep-wake cycle in UWS and healthy controls, we find the spectral slope fits the assumption on arousal in the two-components theory of consciousness, filling the gap in the neural mechanism of arousal, and confirm the relationship between the measurements of complexity (e.g., spectral entropy and Lempel-Ziv complexity) and awareness. These results strongly support the two-components theory of consciousness. Furthermore, the results indicate that it will be important for understanding the neural substrate of consciousness to investigate the interaction between arousal and awareness in future.

## RESOURCE AVAILABILITY

### Lead contact

Further information and requests for resources and reagents should be directed to and will be fulfilled by the lead contact, Yuanqing Li.

### Materials availability

This study did not generate new unique reagents.

### Data and code availability

- The source data for figures have been deposited at Zenodo and are publicly available as of the date of publication.
- The code that supports the findings of this study has been deposited at Zenodo and is publicly available as of the date of publication.
- Any additional information required to reanalyze the data reported in this paper is available from the lead contact upon request.

## ACKNOWLEDGMENTS

We thank all participants for their contribution to this study. This work was supported by the STI 2030 Major Projects (Grant 2022ZD0208900), the Key Research and Development Program of Guangdong Province (Grant 2018B030339001), the Guangdong Basic and Applied Basic Research Foundation (Grants 2024A1515011429 and 2023A1515140100), the National Natural Science Foundation of China (Grant 32371098), the Research Center for Brain Cognition and Human Development, Guangdong (Grant 2024B0303390003), and the Guangzhou Talent Plan (Grant 2024D02J0008).

## AUTHOR CONTRIBUTIONS

Y.Q.L. conceptualized the study. P.M.Q., M.L., and Y.Q.L. designed the methodology. M.L., P.M.Q., J.H.P., and Y.Q.L. conducted the investigation. M.L., Y.B.H., J.H.P., X.Y.B., Y.Z.Y.Y., A.Q.L., Y.H.L., X.C.Y., and D.C. curated the data. M.L. performed the formal analysis. M.L., J.R.H., H.W., and F.W. created the visualizations. Y.Q.L., P.M.Q., and J.H.P. acquired funding. Project administration was managed by P.M.Q. and Y.Q.L. Supervision was provided by Y.Q.L., P.M.Q., and Y.B.H. M.L. and P.M.Q. drafted the manuscript, with critical revisions from P.M.Q., J.H.P., M.L., and Y.Q.L.

## SUPPLEMENTAL INFORMATION INDEX

Figure S1. Etiology-Related Spectral Slope Patterns

Figure S2. Alpha band power distributions of the discovery dataset

Figure S3. Violin plots of alpha band power distribution across sleep stages in the sleep dataset2

Figure S4. Violin plots of spectral slope (30-45Hz) distributions across sleep stages in HC, MCS, and UWS groups of the discovery dataset

Figure S5. Violin plots of spectral slope (30-45Hz) distributions across sleep stages in the sleep dataset2

Table S1. One-way ANCOVA of sleep architecture metrics across the HC, MCS, and UWS groups, controlling for age

Table S2. Summary of sleep stage durations in UWS patients in the Discovery dataset (min)

Table S3. One-way repeated-measures ANOVA (within-subject) of cortical index features across Wake, N3, and REM stages in the HC and MCS groups

Table S4. One-way ANCOVA of cortical index features across Wake, N3, and REM stages in the UWS group, controlling for age

Table S5. One-way ANCOVA of cortical index features across the HC, MCS, and UWS groups during Wake, N3, and REM stages, controlling for age

Table S6. Summary of patients’ clinical status from the Discovery dataset

Table S7. Summary of patients’ clinical status from the BCI task dataset2

## STAR METHODS

### Experimental model and study participant details

#### Sleep dataset1

The discovery dataset comprised 52 participants, including 34 patients with chronic disorders of consciousness (DOC) and 18 HC. Data were collected between February 2023 and January 2024 at the Guangdong Work Injury Rehabilitation Hospital. Among the DOC patients, 16 were diagnosed with UWS (mean age: 51.68 ± 11.05 years; male/female: 9/7), and 18 with MCS (mean age: 51.27 ± 9.63 years; male/female: 15/3). The HC group (mean age: 41.0 ± 9.99 years; male/female: 8/10). Clinical diagnosis was confirmed using the Coma Recovery Scale-Revised (CRS-R), which was administered one week before and on the day of the experiment. In each assessment period, a minimum of two CRS-R assessments were performed conducted by two experienced physicians.

Patients were included according to the following criteria: a diagnosis of UWS or MCS and age between 18 and 70 years (to minimize age dependent effects on sleep). Patients with severe or unstable clinical conditions, pre-existing central nervous system impairments unrelated to disorders of consciousness, or recent use of barbiturates, antipsychotics, or antidepressants within 24 hours were excluded.

Sleep data was collected during overnight using Compumedics Grael Profusion PSG system (version 4.5, build 500, Compumedics Limited). The average recording duration was 8.402 ± 0.54 hours. Recordings included EEG, EOG, and EMG signals. EEG data were recorded according to the international 10–20 system (F3, F4, C3, C4, O1, O2) with a sampling frequency of 256 Hz. EOG and EMG signals were simultaneously captured to provide comprehensive sleep monitoring. Video recordings were synchronized with EEG to annotate key behavioral events (eye-opening moments, rest and agitation for endogenous movements and nursing for exogenous movements). Electrode impedance was maintained below 5 kΩ.

Ethical approval was obtained (approval number: AF/SC-07/2020.03), and informed consent was acquired from legal guardians in accordance with the Declaration of Helsinki. Detailed information can be found in Supplemental Information, Table S6.

#### Sleep dataset2

This independent sleep dataset comprised recordings collected from 37 healthy volunteers (mean age: 26.45 ± 8.11 years, male/female ratio: 20/17), recruited from South China University of Technology between June 2018 and January 2023. Sleep data were collected during overnight polysomnography session using the SynAmps2 amplifier (Compumedics, Neuroscan, Inc., Australia) and a 32-channel EEG cap according to the international 10-20 system. The average recording duration was 7.02 ± 1.34 hours. EEG signals were sampled at 250 Hz, and electrode impedance was maintained below 5 kΩ.

#### BCI task dataset1

The dataset used in this study originates from a brain-computer interface (BCI) experiment conducted with DOC patients performing an audiovisual recognition task. This dataset, previously published in a study investigating the prognostic implications of BCI-based assessments in patients with DOC^77^, comprises EEG recordings from 78 DOC patients, including 45 UWS (mean age: 39.06 ± 14.17 years, male/female: 31/14), and 33 MCS (mean age: 36.24 ± 14.48years, male/female: 20/13). The data were collected during item-selection tasks aimed at detecting conscious awareness at the General Hospital of Guangzhou Military Command of People’s Liberation Army, China, between October 2014 and August 2018. EEG signals were recorded using a 32-channel EEG cap positioned according to the international 10–20 system, with a sampling frequency of 250 Hz. Three BCIs were used, including a hybrid P300 and SSVEP BCI based on photograph stimuli (photograph paradigm), a hybrid P300 and SSVEP BCI based on visual number stimuli (number paradigm), and an audiovisual BCI based on audiovisual number stimuli (audiovisual paradigm). Thirty-five patients participated in the experiment based on the photograph paradigm, 13 in the experiment based on the number paradigm, and 30 in the experiment based on the audiovisual paradigm. In this study, we repurposed this dataset to validate cortical indexes for awareness.

#### BCI task dataset2

The BCI task dataset 2 was collected from an audiovisual multimodal BCI experiment, designed to integrate P300 and auditory steady-state response (ASSR) for assessing awareness in DOC patients. The data were acquired at the Guangdong Work Injury Rehabilitation Hospital, Guangzhou, China, between October 2022 and April 2023, using a SynAmps2 amplifier (Compumedics, Neuroscan, Inc., Australia). Although this dataset was obtained from the same hospital as the sleep dataset1, it was collected from a different cohort of patients with DOC patients.

The dataset included 10 DOC patients, comprising 6 UWS patients (mean age: 49.17 ± 8.45 years, male/female: 4/2) and 4 MCS patients (mean age: 51.75 ± 19.35 years, male/female: 3/1). EEG recordings were conducted over a 5-minute period using a 32-channel EEG cap, following the international 10–20 system, with a sampling frequency of 250 Hz. To ensure high signal fidelity, electrode impedance was maintained below 5 kΩ During each trial, the two facial photographs flashed in a random order and the same side’s earphone played the corresponding sound when each photo flashed. As with the BCI task dataset1, we repurposed this dataset to validate cortical indexes for awareness. Detailed information can be found in Supplemental Information, Table S7.

## Method details

### Sleep Scoring

Polysomnography recordings (EEG, EOG, EMG) were segmented into 30-s epochs for sleep scoring. Given the abnormal sleep patterns in UWS and MCS patients, the standard sleep scoring criteria used in healthy individuals^78^ cannot be applied directly, but have to be adjusted to UWS and MCS patients sleep patterns^13,30,58^. In this study, we applied these adjusted criteria for sleep scoring in UWS and MCS patients.

Specifically, given that UWS and MCS patients may show local intermittent rhythmic delta activity that might be confused with sleep related delta waves, all six EEG electrodes (F3, F4, C3, C4, O1, O2) were utilized for visual sleep scoring to ensure accurate interpretation. Wake state was defined by relatively high EEG frequency, the presence of eye blinks (from video), and high EMG. N1-sleep was scored when the EEG showed a general slowing, accompanied by the disappearance of blinks and decreased muscle activity (rolling eye movements were considered optional indicators). N2-sleep was identified based on the presence of K-complexes and, occasionally, typical sleep spindles (0.5–3 s duration, 20–100 *µ*V amplitude, 11–16 Hz frequency) or prolonged “atypical” slow spindles (10–12 Hz), along with EEG slowing, the emergence of slow-wave activity, and a further reduction in muscle tone, provided that slow waves < 20% of the epoch. N3-sleep was scored when *>* 20% of the epoch consist of slow-wave occurring synchronously across intact cortical regions. Given that EEG amplitude in UWS and MCS patients may be affected by various extracerebral factors, both standard high-amplitude slow waves (*>* 75 *µ*V) and dominant low-voltage slow waves (< 75 *µ*V) were considered indicative of slow-wave sleep (SWS). REM-sleep was defined as fast, low-voltage EEG activity with minimal muscular tone and the presence of muscle atonia and EEG characteristics were prioritized over EOG features^79^.

All sleep data were independently scored by two expert clinical neurophysiologists. Discrepancies in sleep scoring and arousal recognition were reviewed and resolved through consensus. Uncertain epochs were excluded from statistical analyses to maintain data integrity.

### Standard Sleep Architecture Metrics

Total sleep time (TST) was defined as the cumulative duration of all sleep stages (N1, N2, N3, and REM) within the sleep period. Wake time after sleep onset (WASO) was measured as the total duration of wake episodes occurring after the initial sleep onset. Sleep efficiency (SE) is defined as the percentage of total sleep time relative to the total time in bed (TIB, the total duration of the recording period) and was computed as follows:

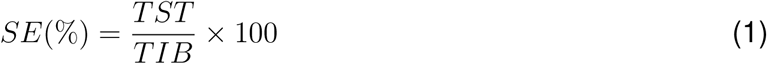

Sleep maintenance efficiency (SME) was defined as the percentage of total sleep time relative to the total sleep period time (SPT):

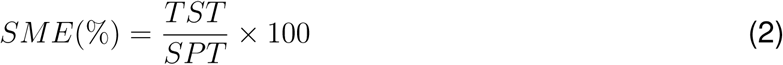

Sleep architecture completeness (SAC) was defined as the proportion of individuals exhibiting a complete sleep architecture, characterized by the presence of all major sleep stages (N1, N2, N3, and REM) during the sleep period. It was computed as:

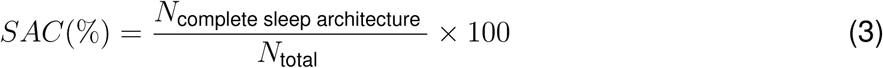

where *N*_complete sleep architecture_ represents the number of individuals experiencing all major sleep stages, and *N*_total_ is the total number of individuals analyzed.

Night wake dominance (NWD) was used to assess the proportion of individuals whose total wake time exceeded total sleep time during the recording period. The calculation was performed as:

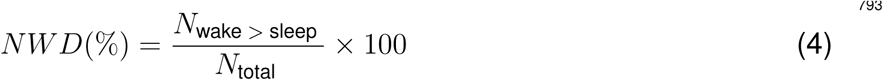

where *N*_wake > sleep_ represents the number of individuals whose total wake time was greater than their TST.

### EEG Preprocessing

A unified preprocessing pipeline was rigorously applied to both sleep EEG datasets (the primary discovery dataset, Sleep dataset1, and the independent sleep dataset, Sleep dataset2) and BCI task datasets (dataset1 and dataset2) using MNE-Python (v1.6.0) integrated with NumPy (v1.22.4) in Python 3.9.6. To ensure methodological consistency across modalities, six electrodes (F3, F4, C3, C4, O1, O2) following the international 10–20 system were selected, encompassing frontal, central, and occipital regions relevant to both sleep and task-related neural dynamics.

The EEG signals were initially processed with a 50 Hz notch filter (4th-order Butterworth) to remove power-line interference, followed by band-pass filtering in the range of 0.5–45 Hz using a finite impulse response (FIR) Hamming window filter. The signals were then re-referenced to the global average of all non-excluded channels to minimize artifacts from electrode displacement during prolonged recordings. Artifact rejection was performed manually by expert clinical neurophysiologists through visual inspection, using criteria such as an amplitude threshold of ±100 *µ*V to identify and exclude segments contaminated by noise. In order to reduce inter-channel variability, z-score normalization was applied independently to each channel. Finally, the signals were down sampled to 100 Hz and segmented into non-overlapping 30-second epochs.

### Feature Extraction

For the discovery dataset (i.e., Sleep dataset 1; comprising UWS, MCS, and HC groups), which was primarily used for statistical analysis, feature extraction was restricted to three sleep stages: Wake-state (high arousal/high awareness), REM-sleep (low arousal/high awareness), and N3-sleep (low arousal/low awareness)as these stages most clearly capture the distinct arousal and awareness profiles pertinent to our study. Notably, previous EEG studies focusing on arousal and consciousness have similarly limited analyses to these three stages^29,32^. Features were initially computed at the epoch level and then averaged for each stage within each subject. For each subject, all three sleep stages (Wake, REM, and N3) included more than 7 epochs (i.e., at least 3.5 minutes of data per stage). Additionally, to obtain a more generalized representation of neural activity, features were further averaged across EEG channels before statistical comparisons. In contrast, for the independent sleep and BCI task datasets (used for sleep staging and consciousness diagnosis, respectively), each epoch was treated as an independent sample.

### Spectral Slope

The spectral slope is a quantitative feature characterizing the power distribution across frequency components, derived from the power spectral density (PSD) analysis of neural signals^25,27,29^. In this study, spectral slope was computed for each channel and epoch following the methods used in previous studies^29,80^. To obtain the spectral slope, we fitted a linear regression line to the power spectrum in log-log space between 0.5 and 45 Hz.

Formally, for a given temporal signal *x*(*t*), the PSD of the signal *x*(*t*) is computed using the “welch” method to ensure robust and smoothed spectral estimation. The PSD is denoted as *P* (*f*) where *f* represents the frequency. The PSD is then transformed into the logarithmic domain:

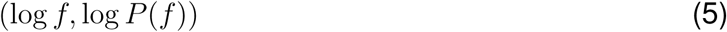

this transformation linearizes the relationship between frequency and power in the log-log space, enabling a more interpretable slope calculation.

A linear regression is performed on the log-transformed PSD values within a predefined frequency range (0.5–45 Hz). The spectral slope is defined as the slope of the regression line:

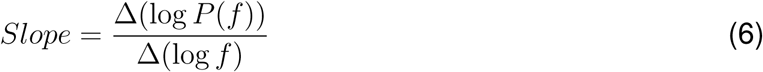

### Spectral Entropy

Spectral entropy(SEp) quantifies the uniformity or peakedness of the EEG power spectral distribution within a specific frequency range^81^. It reflects the degree of randomness or predictability in the frequency components of the signal and provides an indicator of its sinusoidal characteristics. A lower SEp suggests that the signal is dominated by fewer frequency components, with the spectral power concentrated around one or several discrete frequencies, thereby exhibiting a more pronounced sinusoidal nature. In contrast, a higher SEp value indicates a greater number of frequency components with a broader spectral distribution, signifying reduced sinusoidal characteristics and increased complexity of the signal. Building upon established methodologies in neurophysiological signal analysis^20^, we implemented the calculation of spectral entropy within the 0.5-45 Hz bandwidth through three systematic phases for each channel and epoch.

For a continuous signal *x*(*t*), the formula can be defined as follows:

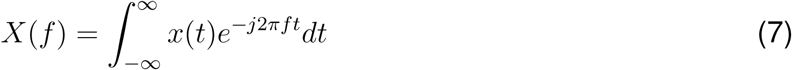

The power spectral density of each frequency component is calculated as follows:

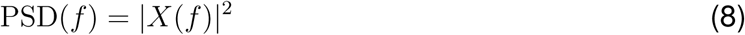

where |*X*(*f*)|^2^ represents the power at frequency, which is the energy of the signal at that frequency. *P* denotes is the relative power of frequency bands *f*:

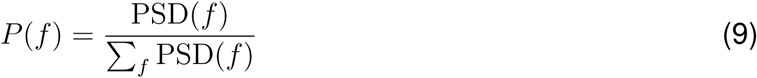

The spectral entropy (SEp) is defined as:

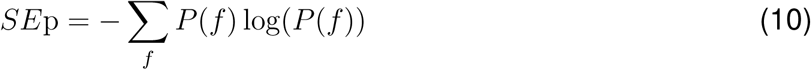

### Lempel-Ziv complexity

Lempel-Ziv Complexity (LZC) was used to quantify the complexity of EEG signals by assessing their temporal dynamics and unpredictability. In this study, LZC was computed for each channel and epoch following the calculation methods used in previous studies^82–84^. We employed a mean symbolic encoding method to compute LZC, where the continuous EEG signal was discretized by mapping it to symbolic representations based on its mean value within a sliding window. To ensure that LZC was computed within the same frequency range as the spectral slope, a 0.5 Hz high-pass filter and a 45 Hz low-pass filter were applied to the EEG signals before computation.

The computation of LZC begins with the preprocessing of the continuous signal *x*(*t*), which is divided into non-overlapping windows of size. For each window *X_i_* = {*x*_1_, *x*_2_, *…, x_L_*}, the mean value of the window *µ_i_* is calculated as:

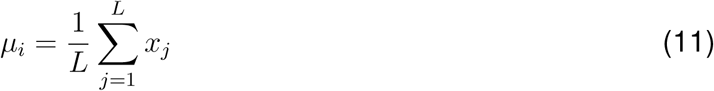

This mean value *µ_i_* is then used to symbolize the window. Specifically, the symbol for each window is determined by comparing the mean value *µ_i_* to predefined thresholds or bins. The EEG signal was transformed into a binary sequence. The Lempel-Ziv parsing algorithm sequentially scans, identifying unique subsequences*S* = {*s*_1_, *s*_2_, *…, s_N_* }. The complexity count *C*(*N*) increases whenever a new, non-redundant subsequence is encountered. To account for sequence length dependency, the raw complexity count *C*(*N*) was normalized by the theoretical upper bound of a random binary sequence:

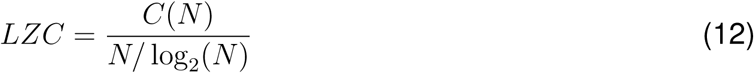

where *N* is the length of the binary sequence. This normalized LZC provides a robust measure of EEG complexity, facilitating comparisons across different conditions and subjects.

### Machine learning-based classification modelling

To evaluate the effectiveness of the identified arousal- and awareness-related features, we constructed a classification framework designed for two tasks: sleep staging and consciousness state diagnosis. The primary motivation was to validate the effectiveness of the cortical index of arousal and awareness.

### Sleep staging

In this study, three separate Support Vector Machine (SVM) classifiers with radial basis function (RBF) kernels were constructed to evaluate the contribution of distinct feature domains to sleep staging. Specifically, the classifiers were trained using arousal-related features (spectral slope), awareness-related features (spectral entropy and Lempel-Ziv complexity), and a combination of both feature types (cortical index of arousal and awareness). All features were extracted from preprocessed EEG data during Wake, N3, and REM stages, as described in the Feature Extraction section.

To account for class imbalance across sleep stages, all classifiers were implemented using scikit-learn (v1.1.3) with class weight = “balanced”. Sleep staging performance was evaluated using leave-one-subject-out cross-validation (LOSO-CV), where each subject was iteratively held out for testing while the remaining data were used for training. Performance metrics included overall classification accuracy (ACC), macro-averaged F1 score (MF1), and Cohen’s Kappa coefficient (K).

### Consciousness state diagnosis

In this study, two separate Support Vector Machine (SVM) classifiers with radial basis function (RBF) kernels were constructed using wake-stage EEG data from the discovery dataset (comprising UWS and MCS groups). One classifier was trained exclusively on the arousal-related feature (spectral slope), while the other utilized awareness-related features (spectral entropy and Lempel-Ziv complexity). Feature extraction procedures followed the same pipeline described in the Feature Extraction section.

To validate the effectiveness of the awareness-related cortical index in distinguishing state of consciousness, both classifiers were independently validated on two external BCI task datasets (dataset 1 and dataset 2), with strict alignment between feature types during training and testing (i.e., the arousal classifier was tested only on spectral slope, and the awareness classifier on spectral entropy and Lempel-Ziv complexity).

For each epoch, the classifiers generated probabilistic outputs indicating the likelihood of the predicted consciousness state (MCS or UWS). To obtain a subject-level diagnosis, a majority-vote strategy was applied across all epochs: if the aggregated probability of being classified as MCS exceeded 0.5, the subject was labeled as MCS; otherwise, UWS. This approach accounted for the dynamic and fluctuating nature of consciousness, particularly in patients with severe brain injuries. The classification performance was assessed using accuracy and the area under the receiver operating characteristic curve (AUC) as primary evaluation metrics.

## Quantification and statistical analysis

All statistical analyses were conducted using SPSS (v27.0, IBM Corp., Armonk, NY, USA). For within-group comparisons across sleep stages (Wake, REM, N3), a one-way repeated-measures ANOVA was applied for the HC and MCS groups, while a one-way ANCOVA with age as a covariate was used for the UWS group due to missing data in certain stages. For between-group comparisons (HC, MCS, UWS), a one-way ANCOVA was performed separately for each sleep stage, also with age as a covariate to control for age-related effects. To further examine etiology-related effects (anoxia vs. no-anoxia), subgroup analyses were conducted. Given the relatively small sample sizes, non-parametric tests were applied (Kruskal–Wallis for within-subgroup stage comparisons and Mann–Whitney U for between-group comparisons). For the MCS no-anoxia subgroup, where sample size was larger and assumptions of normality and sphericity were satisfied, repeated-measures ANOVA was used instead. Post-hoc pairwise comparisons were corrected using the Bonferroni method. Assumptions of sphericity were tested with Mauchly’s test, and the Greenhouse–Geisser correction was applied when violated. Outliers exceeding ±2 SD from the mean were excluded.

