## Supplementary Information Appendix for "Detecting cortical index of arousal in sleeping patients with unresponsive wakefulness syndrome"

### Supporting Information

**This PDF file includes:**

Figures S1 to S5
Tables S1 to S7

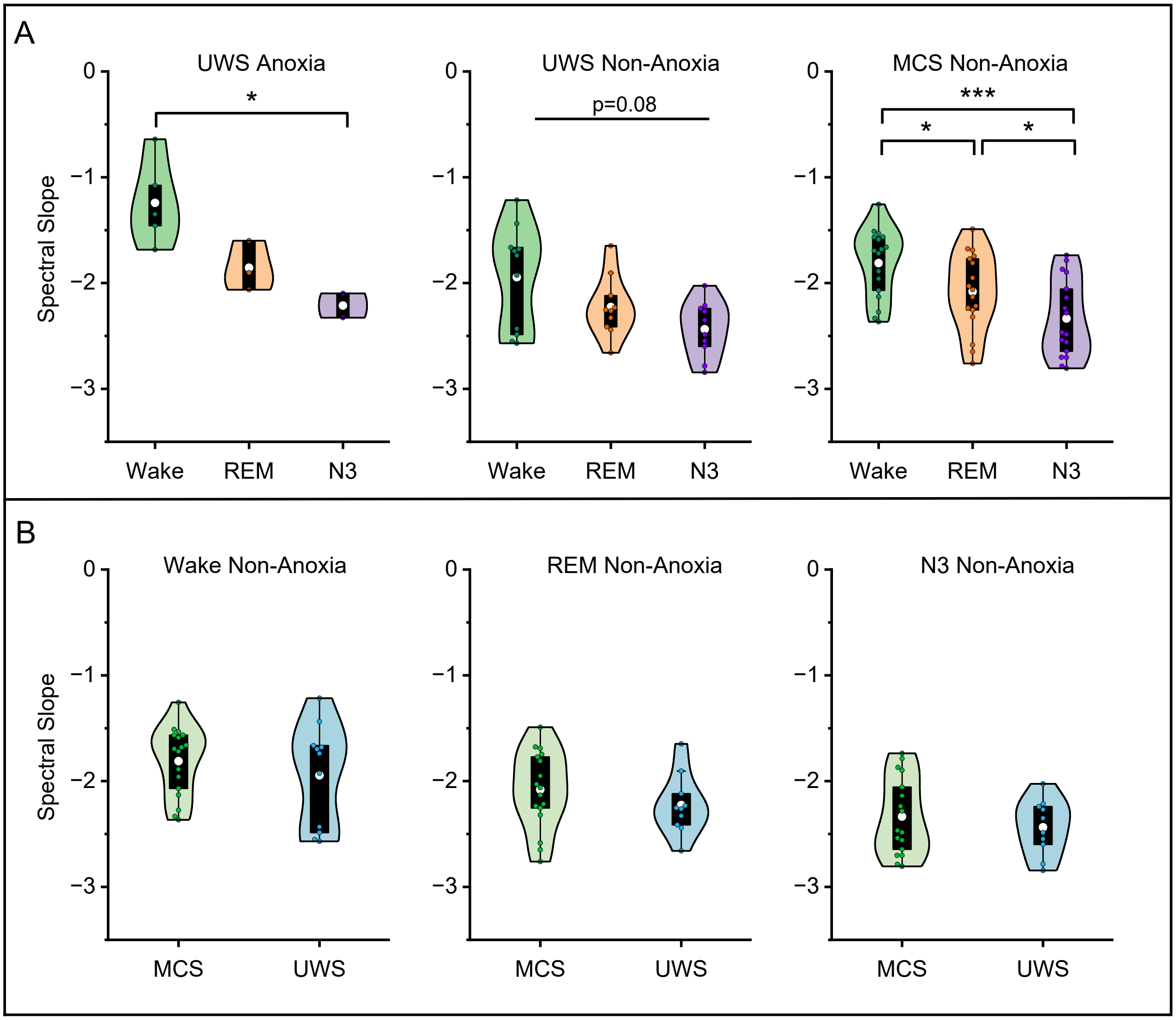


**Figure S1.** Etiology-Related Spectral Slope Patterns. (A) Violin plots of spectral slope distributions across sleep stages (Wake, REM, N3) within UWS anoxia, UWS non-anoxia, and MCS non-anoxia subgroups. (B) Violin plots of spectral slope distributions for between-group comparisons (MCS vs. UWS) under non-anoxia conditions for each sleep stage. The white dot indicates the mean, and the box boundaries represent the upper and lower quartiles.$*p < 0.05, **p < 0.01, ***p < 0.001$.

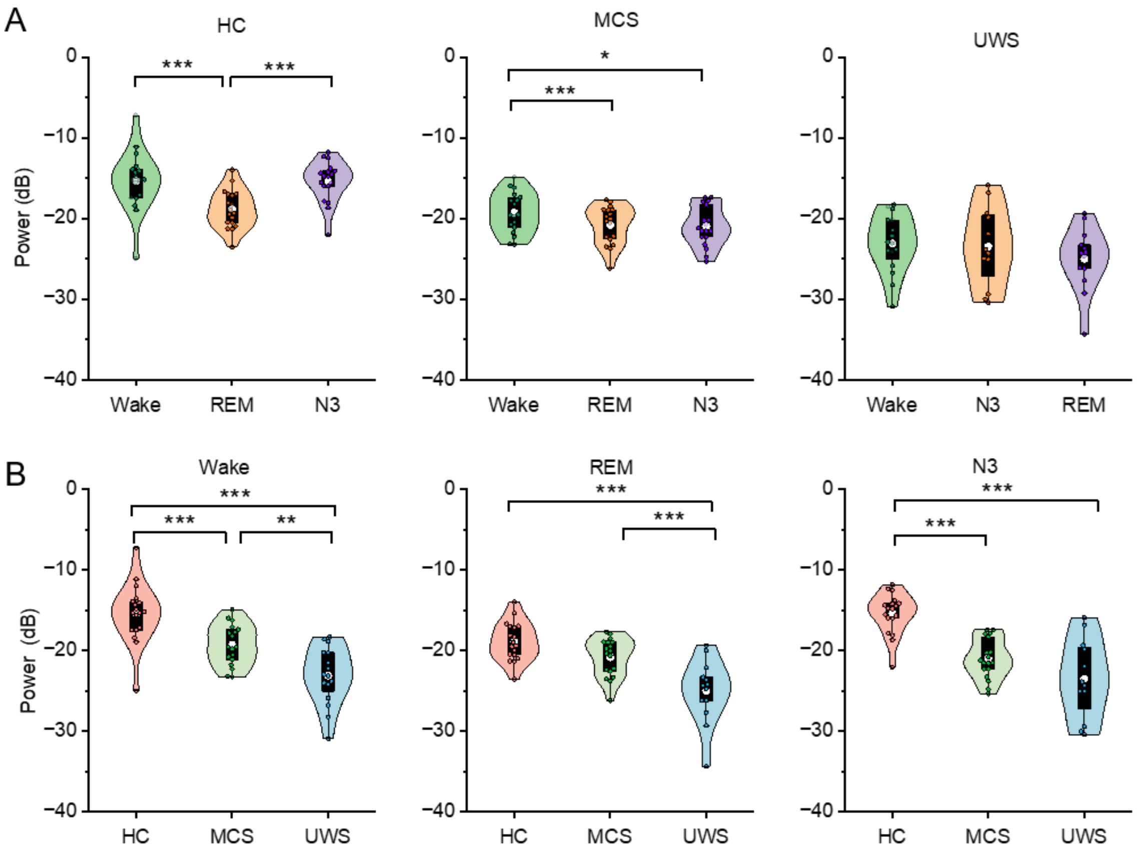


**Figure S2.** Alpha band power distributions of the discovery dataset. (A)Violin plots of alpha power distributions for each sleep stage within HC, MCS, and UWS groups. (B) Violin plots of alpha power distributions for individual sleep stages across HC, MCS, and UWS groups. The white dot indicates the mean, and the box boundaries represent the upper and lower quartiles.$*p < 0.05, **p < 0.01, ***p < 0.001$.

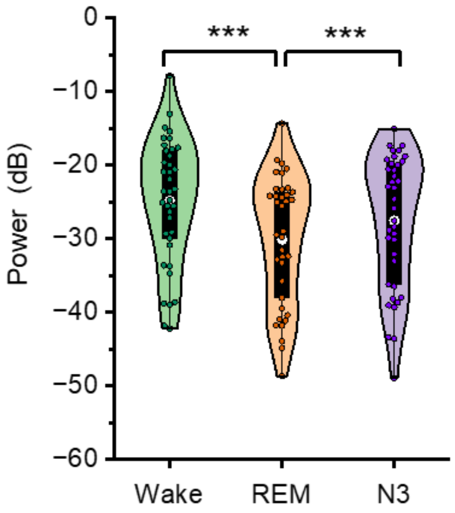


**Figure S3.** Violin plots of alpha band power distribution across sleep stages in the sleep dataset2. The white dot indicates the mean, and the box boundaries represent the upper and lower quartiles.$*p < 0.05, **p < 0.01, ***p < 0.001$.

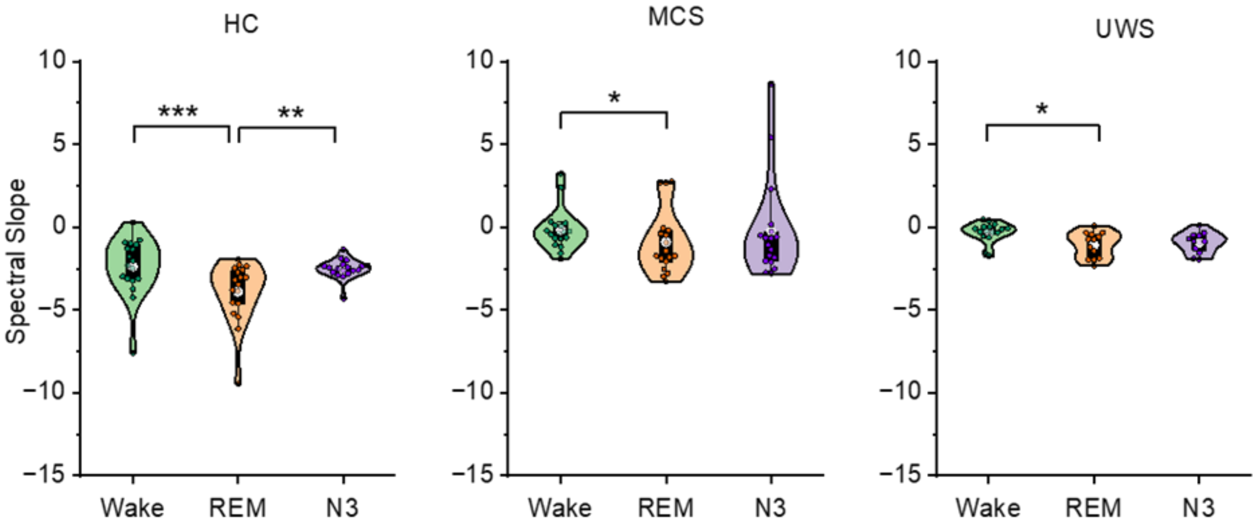


**Figure S4.** Violin plots of spectral slope (30-45Hz) distributions across sleep stages in HC, MCS, and UWS groups of the discovery dataset. The white dot indicates the mean, and the box boundaries represent the upper and lower quartiles.$*p < 0.05, **p < 0.01, ***p < 0.001$.

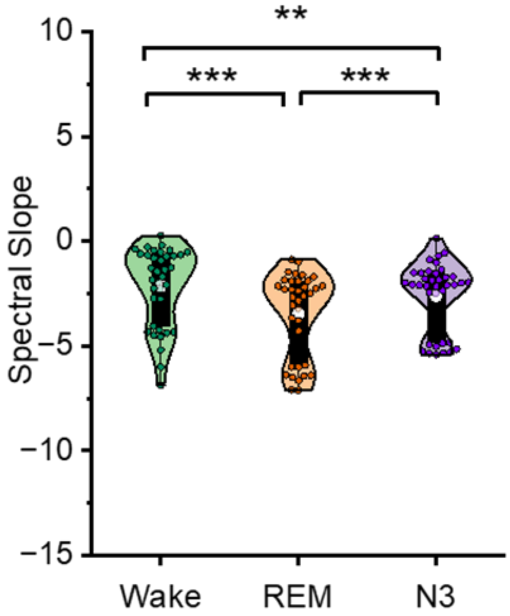


**Figure S5.** Violin plots of spectral slope (30-45Hz) distributions across sleep stages in the sleep dataset2. The white dot indicates the mean, and the box boundaries represent the upper and lower quartiles.$*p < 0.05, **p < 0.01, ***p < 0.001.$

**Table S1. One-way ANCOVA of sleep architecture metrics across the HC, MCS, and UWS groups, controlling for age.**

| Sleep Stage | Main Effect (Group: F(df), p, $\eta_p^2$) | Covariate (Age: F(df), p, $\eta_p^2$) | Post hoc comparisons (Bonferroni-corrected) | | | | |
|---|---|---|---|---|---|---|---|
| | | | Group Comparison | MD | p | Cohen's d | [95% CI] |
| TST | F(2,47) = 13.971, p $<$ 0.001***, $\eta_p^2$ = 0.373 | F(1,47) = 0.533, p = 0.469, $\eta_p^2$ = 0.011 | HC vs. MCS | 90.674 | 0.03* | 0.985 | [0.041, 1.930] |
| | | | HC vs. UWS | 99.178 | $<$0.001*** | 1.993 | [0.924, 3.061] |
| | | | MCS vs. UWS | 16.174 | 0.016* | 1.007 | [0.116, 1.899] |
| WASO | F(2,48) = 14.401, p $<$ 0.001***, $\eta_p^2$ = 0.375 | F(1,48) = 0.530, p = 0.470, $\eta_p^2$ = 0.011 | HC vs. MCS | -106.341 | 0.003** | -1.263 | [-2.219, -0.307] |
| | | | HC vs. UWS | -168.425 | $<$0.001*** | -2.000 | [-3.059, -0.942] |
| | | | MCS vs. UWS | -62.084 | 0.111 | -0.737 | [-1.610, 0.135] |
| SE | F(2,47) = 13.728, p $<$ 0.001***, $\eta_p^2$ = 0.369 | F(1,47) = 0.161, p = 0.690, $\eta_p^2$ = 0.003 | HC vs. MCS | 19.256 | 0.022* | 1.031 | [0.083, 1.978] |
| | | | HC vs. UWS | 36.977 | $<$0.001*** | 1.979 | [0.912, 3.046] |
| | | | MCS vs. UWS | 17.721 | 0.025* | 0.949 | [0.061, 1.836] |
| SME | F(2,48) = 17.104, p $<$ 0.001***, $\eta_p^2$ = 0.416 | F(1,48) = 0.078, p = 0.781, $\eta_p^2$ = 0.002 | HC vs. MCS | 20.233 | 0.01* | 1.125 | [0.181, 2.070] |
| | | | HC vs. UWS | 39.367 | $<$0.001*** | 2.189 | [1.107, 3.272] |
| | | | MCS vs. UWS | 19.134 | 0.01* | 1.064 | [0.170, 1.958] |

**Note.** Abbreviations used in this table are defined as follows: TST, total sleep time; WASO, wake time after sleep onset; SE, sleep efficiency; SME, sleep maintenance efficiency. Notably, one HC subject with both TST and SE values exceeding 2 standard deviations below the group mean was excluded from analyses (TST = 103.5 min; group mean ± SD = 384.44 ± 84.00 min; SE = 20.51%; group mean ± SD = 77.94 ± 17.33%). $^{*}p < 0.05$, $^{**}p < 0.01$, $^{***}p < 0.001$.

**Table S2. Summary of sleep stage durations in UWS patients in the Discovery dataset (min).**

| Patient | N1 (min) | N2 (min) | N3 (min) | REM (min) |
|---|---|---|---|---|
| UWS1 | 7.5 | 121.5 | 31.5 | 121 |
| UWS2 | 1.5 | 11 | 11 | 25 |
| UWS3 | 21 | 203.5 | 33.5 | 110 |
| UWS4 | 0.5 | 26.5 | 29.5 | 13 |
| UWS5 | 0 | 1.5 | 0 | 35.5 |
| UWS6 | 10 | 199.5 | 75.5 | 15.5 |
| UWS7 | 2.5 | 210.5 | 78 | 39.5 |
| UWS8 | 5.5 | 64.5 | 86 | 148 |
| UWS9 | 38 | 7 | 0 | 0 |
| UWS10 | 15.5 | 194.5 | 0 | 37 |
| UWS11 | 9 | 114.5 | 126 | 38 |
| UWS12 | 23.5 | 130.5 | 100 | 0 |
| UWS13 | 39.5 | 117.5 | 0 | 114.5 |
| UWS14 | 6.5 | 138 | 26.5 | 0 |
| UWS15 | 59.5 | 142.5 | 14 | 145 |
| UWS16 | 17.5 | 221 | 15 | 61 |

**Table S3. One-way repeated-measures ANOVA (within-subject) of cortical index features across Wake, N3, and REM stages in the HC and MCS groups.**

| Feature | Group | Main effect (Stage: F(df), p, $\eta_p^2$) | Post hoc comparisons (Bonferroni-corrected) | | | | |
|---|---|---|---|---|---|---|---|
| | | | Stage Comparison | MD | p | Cohen's d | [95% CI] |
| Spectral slope | HC | F(1.404, 23.860) = 242.340, p < 0.001***, $\eta_p^2$ = 0.934 | Wake vs. REM | 0.369 | <0.001*** | 1.596 | [0.642, 2.550] |
| | | | Wake vs. N3 | 1.316 | <0.001*** | 5.691 | [3.164, 8.218] |
| | | | REM vs. N3 | 0.947 | <0.001*** | 4.095 | [2.219, 5.971] |
| | MCS | F(2,34) = 27.661, p < 0.001***, $\eta_p^2$ = 0.619 | Wake vs. REM | 0.293 | <0.001*** | 0.852 | [0.203, 1.502] |
| | | | Wake vs. N3 | 0.537 | <0.001*** | 1.560 | [0.697, 2.424] |
| | | | REM vs. N3 | 0.244 | 0.011* | 0.708 | [0.092, 1.324] |
| Spectral entropy | HC | F(1.283, 21.803) = 66.073, p < 0.001***, $\eta_p^2$ = 0.795 | Wake vs. REM | 0.003 | >0.999 | 0.005 | [-0.564, 0.575] |
| | | | Wake vs. N3 | 1.078 | <0.001*** | 2.247 | [1.119, 3.375] |
| | | | REM vs. N3 | 1.076 | <0.001*** | 2.241 | [1.116, 3.367] |
| | MCS | F(1.490, 25.328) = 21.773, p < 0.001***, $\eta_p^2$ = 0.562 | Wake vs. REM | 0.050 | >0.999 | 0.115 | [-0.554, 0.784] |
| | | | Wake vs. N3 | 0.687 | <0.001*** | 1.581 | [0.630, 2.531] |
| | | | REM vs. N3 | 0.637 | <0.001*** | 1.466 | [0.550, 2.382] |
| Lempel-Ziv complexity | HC | F(1.455, 24.743) = 127.414, p < 0.001***, $\eta_p^2$ = 0.882 | Wake vs. REM | 0.051 | 0.086 | 0.684 | [0.032, 1.336] |
| | | | Wake vs. N3 | 0.259 | <0.001*** | 3.472 | [1.862, 5.082] |
| | | | REM vs. N3 | 0.208 | <0.001*** | 2.788 | [1.449, 4.126] |
| | MCS | F(2,34) = 27.550, p < 0.001***, $\eta_p^2$ = 0.618 | Wake vs. REM | 0.023 | 0.326 | 0.305 | [-0.317, 0.928] |
| | | | Wake vs. N3 | 0.127 | <0.001*** | 1.686 | [0.738, 2.634] |
| | | | REM vs. N3 | 0.104 | <0.001*** | 1.380 | [0.529, 2.232] |
| Alpha power | HC | F(1.458, 24.789) = 20.006, p < 0.001***, $\eta_p^2$ = 0.541 | Wake vs. REM | 3.472 | <0.001*** | 1.187 | [0.434, 1.939] |
| | | | Wake vs. N3 | 0.009 | >0.999 | 0.003 | [-0.545,0.551] |
| | | | REM vs. N3 | -3.464 | <0.001*** | -1.184 | [-1.935,-0.432] |
| | MCS | F(1.372, 23.320) = 9.420, p = 0.003**, $\eta_p^2$ = 0.357 | Wake vs. REM | 1.546 | <0.001*** | 0.630 | [0.100, 1.161] |
| | | | Wake vs. N3 | 1.694 | 0.017* | 0.691 | [0.146, 1.236] |
| | | | REM vs. N3 | 0.149 | >0.999 | 0.061 | [-0.392, 0.514] |

**Note.** $^{*}p < 0.05$, $^{**}p < 0.01$, $^{***}p < 0.001$.

**Table S4. One-way ANCOVA of cortical index features across Wake, N3, and REM stages in the UWS group, controlling for age.**

| Feature | Main effect (Stage: F(df), p, $\eta_p^2$) | Covariate (Age: F(df), p, $\eta_p^2$) | Post hoc comparisons (Bonferroni-corrected) | | | | |
|---|---|---|---|---|---|---|---|
| | | | Stage Comparison | MD | p | Cohen's d | [95% CI] |
| Spectral slope | F(2, 37) = 9.259, p < 0.001***, $\eta_p^2$ = 0.334 | F(1, 37) = 0.497, p = 0.485, $\eta_p^2$ = 0.013 | Wake vs. REM | 0.409 | 0.039* | 0.977 | [–0.003, 1.957] |
| | | | Wake vs. N3 | 0.674 | <0.001*** | 1.611 | [0.544, 2.677] |
| | | | REM vs. N3 | 0.265 | 0.368 | 0.634 | [–0.389, 1.656] |
| Spectral entropy | F(2, 37) = 1.085, p = 0.348, $\eta_p^2$ = 0.055 | F(1, 37) = 3.804, p = 0.059, $\eta_p^2$ = 0.093 | Wake vs. REM | –0.058 | >0.999 | –0.102 | [–1.040, 0.837] |
| | | | Wake vs. N3 | 0.260 | 0.733 | 0.452 | [–0.515, 1.419] |
| | | | REM vs. N3 | 0.318 | 0.527 | 0.553 | [–0.465, 1.572] |
| Lempel-Ziv complexity | F(2, 37) = 2.481, p = 0.097, $\eta_p^2$ = 0.118 | F(1, 37) = 0.799, p = 0.377, $\eta_p^2$ = 0.021 | Wake vs. REM | 0.026 | >0.999 | 0.278 | [–0.663, 1.219] |
| | | | Wake vs. N3 | 0.078 | 0.099 | 0.845 | [–0.144, 1.834] |
| | | | REM vs. N3 | 0.052 | 0.496 | 0.568 | [–0.451, 1.587] |
| Alpha power | F(2, 37) = 0.887, p = 0.420, $\eta_p^2$ = 0.046 | F(1, 37) = 0.703, p = 0.407, $\eta_p^2$ = 0.019 | Wake vs. REM | 1.984 | 0.624 | 0.479 | [–0.469, 1.427] |
| | | | Wake vs. N3 | 0.369 | >0.999 | 0.089 | [–0.869, 1.047] |
| | | | REM vs. N3 | –1.615 | >0.999 | –0.390 | [–1.402, 0.622] |

**Note.** $^*p < 0.05$, $^{**}p < 0.01$, $^{***}p < 0.001$.

**Table S5. One-way ANCOVA of cortical index features across the HC, MCS, and UWS groups during Wake, N3, and REM stages, controlling for age.**

| Feature | Sleep stage | Main effect (Group: F(df), p, $\eta_p^2$) | Covariate (Age: F(df), p, $\eta_p^2$) | Post hoc comparisons (Bonferroni-corrected) | | | | |
|---|---|---|---|---|---|---|---|---|
| | | | | Group Comparison | MD | p | Cohen's d | [95% CI] |
| Spectral slope | Wake | F(2,48) = 2.211, p = 0.121, $\eta_p^2$ = 0.084 | F(1,48) = 0.995, p = 0.324, $\eta_p^2$ = 0.020 | HC vs. MCS | 0.298 | 0.134 | 0.749 | [-0.171, 1.669] |
| | | | | HC vs. UWS | 0.224 | 0.418 | 0.563 | [-0.377, 1.504] |
| | | | | MCS vs. UWS | -0.074 | >0.999 | -0.186 | [-1.040, 0.668] |
| | REM | F(2,45) = 4.020, p = 0.025*, $\eta_p^2$ = 0.152 | F(1,45) = 0.058, p = 0.810, $\eta_p^2$ = 0.001 | HC vs. MCS | 0.268 | 0.059 | 0.888 | [-0.054, 1.830] |
| | | | | HC vs. UWS | 0.316 | 0.040* | 1.407 | [0, 2.095] |
| | | | | MCS vs. UWS | 0.048 | >0.999 | 0.159 | [-0.749, 1.068] |
| | N3 | F(2,44) = 8.271, p < 0.001**, $\eta_p^2$ = 0.273 | F(1,44) = 0.331, p = 0.568, $\eta_p^2$ = 0.007 | HC vs. MCS | -0.401 | <0.001*** | -1.426 | [-2.406, -0.446] |
| | | | | HC vs. UWS | -0.336 | 0.014* | -1.194 | [-1.194, -2.241] |
| | | | | MCS vs. UWS | 0.065 | >0.999 | 0.232 | [-0.698, 1.162] |
| Spectral entropy | Wake | F(2,48) = 26.139, p < 0.001***, $\eta_p^2$ = 0.521 | F(1,48) = 1.174, p = 0.284, $\eta_p^2$ = 0.024 | HC vs. MCS | 0.814 | <0.001*** | 1.586 | [0.600, 2.572] |
| | | | | HC vs. UWS | 1.389 | <0.001*** | 2.706 | [1.551, 3.861] |
| | | | | MCS vs. UWS | 0.575 | 0.006** | 1.120 | [0.222, 2.018] |
| | REM | F(2,45) = 22.422, p < 0.001***, $\eta_p^2$ = 0.499 | F(1,45) = 2.504, p = 0.121, $\eta_p^2$ = 0.053 | HC vs. MCS | 0.906 | <0.001*** | 1.763 | [0.740, 2.786] |
| | | | | HC vs. UWS | 1.369 | <0.001*** | 2.663 | [1.435, 3.891] |
| | | | | MCS vs. UWS | 0.463 | 0.053 | 0.900 | [-0.038, 1.838] |
| | N3 | F(2,44) = 8.008, p = 0.001***, $\eta_p^2$ = 0.267 | F(1,44) = 6.153, p = 0.017*, $\eta_p^2$ = 0.123 | HC vs. MCS | 0.497 | 0.008** | 1.153 | [0.198, 2.107] |
| | | | | HC vs. UWS | 0.648 | 0.002** | 1.502 | [0.427, 2.576] |
| | | | | MCS vs. UWS | 0.150 | >0.999 | 0.349 | [-0.584, 1.281] |
| Lempel-Ziv complexity | Wake | F(2,48) = 27.608, p < 0.001***, $\eta_p^2$ = 0.535 | F(1,48) = 0.350, p = 0.557, $\eta_p^2$ = 0.007 | HC vs. MCS | 0.166 | <0.001*** | 1.877 | [0.859, 2.895] |
| | | | | HC vs. UWS | 0.243 | <0.001*** | 2.745 | [1.585, 3.906] |
| | | | | MCS vs. UWS | 0.077 | 0.045* | 0.868 | [-0.012, 1.748] |
| | REM | F(2,45) = 25.246, p < 0.001***, $\eta_p^2$ = 0.529 | F(1,45) = 1.924, p = 0.172, $\eta_p^2$ = 0.041 | HC vs. MCS | 0.148 | <0.001*** | 1.836 | [0.804, 2.867] |
| | | | | HC vs. UWS | 0.228 | < 0.001*** | 2.836 | [1.582, 4.091] |
| | | | | MCS vs. UWS | 0.081 | 0.026* | 1.001 | [0.056, 1.946] |
| | N3 | F(2,44) = 4.154, p = 0.022*, $\eta_p^2$ = 0.159 | F(1,44) = 4.320, p = 0.044*, $\eta_p^2$ = 0.089 | HC vs. MCS | 0.046 | 0.178 | 0.703 | [-0.220, 1.627] |
| | | | | HC vs. UWS | 0.075 | 0.021* | 1.136 | [0.094, 2.179] |
| | | | | MCS vs. UWS | 0.029 | 0.755 | 0.433 | [-0.502, 1.368] |
| Alpha power | Wake | F(2,48) = 26.848, p < 0.001***, $\eta_p^2$ = 0.528 | F(1,48) = 4.469, p = 0.040*, $\eta_p^2$ = 0.085 | HC vs. MCS | 4.869 | <0.001*** | 1.531 | [0.550, 2.511] |
| | | | | HC vs. UWS | 8.732 | <0.001*** | 2.745 | [1.585, 3.906] |
| | | | | MCS vs. UWS | 3.863 | 0.003** | 1.215 | [0.308, 2.121] |
| | REM | F(2,45) = 17.463, p < 0.001***, $\eta_p^2$ = 0.437 | F(1,45) = 1.346, p = 0.252, $\eta_p^2$ = 0.029 | HC vs. MCS | 2.487 | 0.067 | 0.868 | [-0.073, 1.808] |
| | | | | HC vs. UWS | 6.770 | <0.001*** | 2.362 | [1.177, 3.547] |
| | | | | MCS vs. UWS | 4.283 | <0.001*** | 1.494 | [0.506, 2.483] |
| | N3 | F(2,44) = 20.700, p < 0.001***, $\eta_p^2$ = 0.485 | F(1,44) = 0.035, p = 0.853, $\eta_p^2$ = 0.001 | HC vs. MCS | 5.677 | <0.001*** | 1.722 | [0.709, 2.735] |
| | | | | HC vs. UWS | 8.188 | <0.001*** | 2.484 | [1.288, 3.680] |
| | | | | MCS vs. UWS | 2.511 | 0.141 | 0.762 | [-0.188, 1.711] |

**Note.** $^{*}p < 0.05$, $^{**}p < 0.01$, $^{***}p < 0.001$. MD = Mean Difference; CI = Confidence Interval.

**Table S6. Summary of patients' clinical status from the Discovery dataset.**

| Patient | Age (years) | Gender | Aetiology | Time since injury (months) | Before experiment | | Follow-up Status |
|---|---|---|---|---|---|---|---|
| | | | | | CRS-R score (subscores) | Diagnosis | |
| UWS1 | 50 | F | CVD | 8.5 | 6 (1-1-1-1-0-2) | UWS | LTFU |
| UWS2 | 69 | M | CVD | 18 | 6 (1-1-1-1-0-2) | UWS | FUC |
| UWS3 | 34 | M | TBI | 12 | 4 (1-0-1-0-0-2) | UWS | FUC |
| UWS4 | 52 | F | CVD | 8 | 5 (1-0-2-0-0-2) | UWS | FUC |
| UWS5 | 52 | F | TBI | 10 | 5 (1-1-1-0-0-2) | UWS | LTFU |
| UWS6 | 48 | M | CVD | 24 | 8 (2-1-2-1-0-2) | UWS | FUC |
| UWS7 | 65 | M | TBI | 2 | 3 (1-0-0-0-0-2) | UWS | FUC |
| UWS8 | 37 | M | TBI | 1 | 5 (0-0-2-1-0-2) | UWS | LTFU |
| UWS9 | 50 | F | ABI | 6 | 6 (1-1-2-0-0-2) | UWS | FUC |
| UWS10 | 58 | F | ABI | 25 | 6 (1-0-2-1-0-2) | UWS | FUC |
| UWS11 | 59 | M | CVD | 2 | 7 (1-1-2-1-0-2) | UWS | FUC |
| UWS12 | 59 | F | TBI | 8 | 5 (1-1-0-1-0-2) | UWS | FUC |
| UWS13 | 49 | M | ABI | 21 | 5 (1-0-1-1-0-2) | UWS | FUC |
| UWS14 | 28 | M | ABI | 3 | 4 (0-0-1-1-0-2) | UWS | FUC |
| UWS15 | 59 | M | TBI | 1 | 8 (2-1-2-1-0-2) | UWS | FUC |
| UWS16 | 58 | F | ABI | 24 | 6 (1-0-2-1-0-2) | UWS | FUC |
| MCS1 | 65 | M | CVD | 6 | 7 (1-3-1-0-0-2) | MCS | FUC |
| MCS2 | 62 | M | CVD | 3 | 8 (1-1-5-0-0-1) | MCS | FUC |
| MCS3 | 54 | M | CVD | 5 | 7 (1-2-2-0-0-2) | MCS | FUC |
| MCS4 | 46 | M | ABI | 3 | 12 (2-3-3-1-1-2) | MCS | FUC |
| MCS5 | 63 | M | TBI | 5 | 9 (0-3-3-1-0-2) | MCS | FUC |
| MCS6 | 47 | M | TBI | 3 | 10 (2-3-2-1-0-2) | MCS | FUC |
| MCS7 | 50 | M | CVD | 12 | 8 (1-1-3-1-0-2) | MCS | FUC |
| MCS8 | 34 | M | TBI | 13 | 9 (1-3-2-1-0-2) | MCS | FUC |
| MCS9 | 56 | F | CVD | 4 | 8 (1-1-3-0-1-2) | MCS | LTFU |
| MCS10 | 54 | M | CVD | 6 | 7 (1-2-2-0-0-2) | MCS | FUC |
| MCS11 | 41 | M | CVD | 5 | 8 (1-0-5-1-0-1) | MCS | LTFU |
| MCS12 | 54 | M | CVD | 7 | 9 (1-3-3-0-0-2) | MCS | FUC |
| MCS13 | 66 | M | TBI | 9 | 10 (1-1-5-1-0-2) | MCS | FUC |
| MCS14 | 50 | F | TBI | 24 | 14 (2-3-5-2-0-2) | MCS | FUC |
| MCS15 | 57 | F | TBI | 4 | 13 (2-2-5-1-0-3) | MCS | FUC |
| MCS16 | 49 | M | TBI | 10 | 7 (1-0-3-1-0-2) | MCS | FUC |
| MCS17 | 39 | M | TBI | 216 | 10 (3-3-1-1-0-2) | MCS | FUC |
| MCS18 | 36 | M | CVD | 12 | 7 (1-0-3-1-0-2) | MCS | FUC |

**Note.** Abbreviations used in this table are defined as follows: UWS, unresponsive wakefulness syndrome; MCS, minimally conscious state; CRS-R, Coma Recovery Scale–Revised; ABI, anoxic brain injury; CVD, cerebrovascular disease; TBI, traumatic brain injury; M, male; F, female; FUC , follow-up completed; LTFU, lost to follow-up. The CRS-R subscales are presented in the following order: auditory, visual, motor, oromotor, communication, and arousal functions.

**Table S7. Summary of patients' clinical status from the BCI task dataset2.**

| Patient | Age (years) | Gender | Etiology | Time since injury (months) | Before experiment | |
|---|---|---|---|---|---|---|
| | | | | | CRS-R score (subscores) | Diagnosis |
| UWS1 | 51 | M | ABI | 10 | 5 (0-0-2-1-0-2) | UWS |
| UWS2 | 36 | F | ABI | 4 | 6 (1-0-2-1-0-2) | UWS |
| UWS3 | 57 | M | TBI | 8 | 4 (1-0-0-1-0-2) | UWS |
| UWS4 | 59 | M | TBI | 12 | 8 (2-1-2-1-0-2) | UWS |
| UWS5 | 47 | F | ABI | 9 | 5 (1-1-0-1-0-2) | UWS |
| UWS6 | 45 | M | TBI | 13 | 7 (1-1-2-1-0-2) | UWS |
| MCS1 | 57 | M | ABI | 6 | 8 (3-0-2-1-1-2) | MCS |
| MCS2 | 24 | M | TBI | 15 | 12 (1-3-5-1-0-2) | MCS |
| MCS3 | 69 | F | ABI | 9 | 11 (2-3-2-2-0-2) | MCS |
| MCS4 | 57 | M | ABI | 6 | 9 (3-0-2-1-1-2) | MCS |

**Note.** Abbreviations used in this table are defined as follows: UWS, unresponsive wakefulness syndrome; MCS, minimally conscious state; CRS-R, Coma Recovery Scale–Revised; ABI, anoxic brain injury; TBI, traumatic brain injury; M, male; F, female; The CRS-R subscales are presented in the following order: auditory, visual, motor, oromotor, communication, and arousal functions.
